# The Role of Chloride Ions in Serotonin Transport

**DOI:** 10.1101/2025.05.20.654092

**Authors:** Jiahui Huang, Annika Backer, Stacy Uchendu, Bethlehem Bekele, Chan Li, Qingyang Chen, Esam Orabi, Robyn Stix, Jasper Shide, Yuan-Wei Zhang, Gary Rudnick, Eva Hellsberg, Lucy R. Forrest

## Abstract

The human serotonin (5-HT^+^) transporter SERT facilitates 5-HT^+^ transport into cells by coupling it to Na^+^ symport and K^+^ antiport. Although extracellular Cl^−^ is also essential for transport, Cl^−^co-transport has been disputed, raising questions about the role of Cl^−^ ions and why they are required. We show that Cl^−^ gradients do not impact 5-HT^+^ accumulation, indicating that Cl⁻ does not provide a driving force for uptake and arguing against stoichiometric Cl⁻ symport. The presence of Cl^−^ had only a small effect on Na^+^-mediated cytoplasmic pathway closure but markedly reduced the accessibility of residues in the extracellular pathway, consistent with modulation of the outward-facing states. Simulations illustrate that Cl^−^ interacts strongly with a bound Na^+^ ion and stabilizes helix packing on the extracellular side. We propose that Cl⁻ acts as an essential architectural cofactor by enhancing Na^+^ affinity and interactions between helices, thereby facilitating transport-related conformational transitions.

**Significance statement:** The serotonin transporter (SERT) clears serotonin from synapses and is the target of widely used antidepressants. Although chloride ions are known to be required for SERT function, whether chloride is transported together with serotonin has remained unresolved for decades. We show that chloride gradients do not drive serotonin uptake, arguing against chloride symport. Instead, chloride stabilizes specific conformations of the transporter by strengthening sodium binding and extracellular helix packing. These findings redefine chloride not as a transported substrate, but as an architectural cofactor that facilitates conformational transitions required for transport. This reinterpretation reshapes our understanding of ion dependence in SERT and suggests a broader principle: ion requirements in related transporters may reflect structural stabilization rather than obligatory symport.

## Introduction

Transporters can couple the energy provided by a transmembrane ion concentration difference to the energetically unfavorable accumulation of the transported substrate. This ability depends on the binding of required ligands to control the conformational changes that physically move the substrate and co-transported ions across the membrane. Serotonin transporter (SERT, SLC6A4), a member of the NSS (<u>N</u>eurotransmitter:<u>S</u>odium:<u>S</u>ymporter) family, catalyzes Na^+^ and Cl^−^-dependent accumulation of serotonin (5-hydroxytryptamine, 5-HT^+^) within serotonergic neurons, platelets and other cells, driven by the movement of Na^+^ into the cell and K^+^ out of the cell (1). Other members of the NSS family include transporters for the neurotransmitters dopamine, norepinephrine, glycine, and GABA (γ-aminobutyric acid).

Na^+^ stabilizes SERT in an outward-open conformation (2, 3) and the subsequent binding of Cl^−^and 5-HT^+^ from the cell exterior allows the transformation of SERT to an inward-open conformation (4, 5) from which 5-HT^+^ and Na^+^ dissociate, thereby coupling their transmembrane movement (6). To return SERT to the outward-open state, cytoplasmic K^+^ binds (7–9), replacing Na^+^ and allowing the major conformational reorientation; an additional ion of unknown type may also be translocated to preserve the reported electroneutrality of the transport cycle (7). These conformational changes involve movement of transmembrane (TM) helices in the so-called bundle (TM helices 1, 2, 6, and 7; **Fig. 1A, B**) as well as TM5 in the scaffold region (TM helices 3-5 and 8-12), as indicated by the measured accessibility of cysteine residues placed into the cytoplasmic and extracellular pathways at positions accessible to aqueous reagents in one conformation and less accessible in another (10, 11) (with additional, more distributed changes detected by HDX-MS in TM1, extracellular loops, and TM12 (12)).

**Fig. 1.**
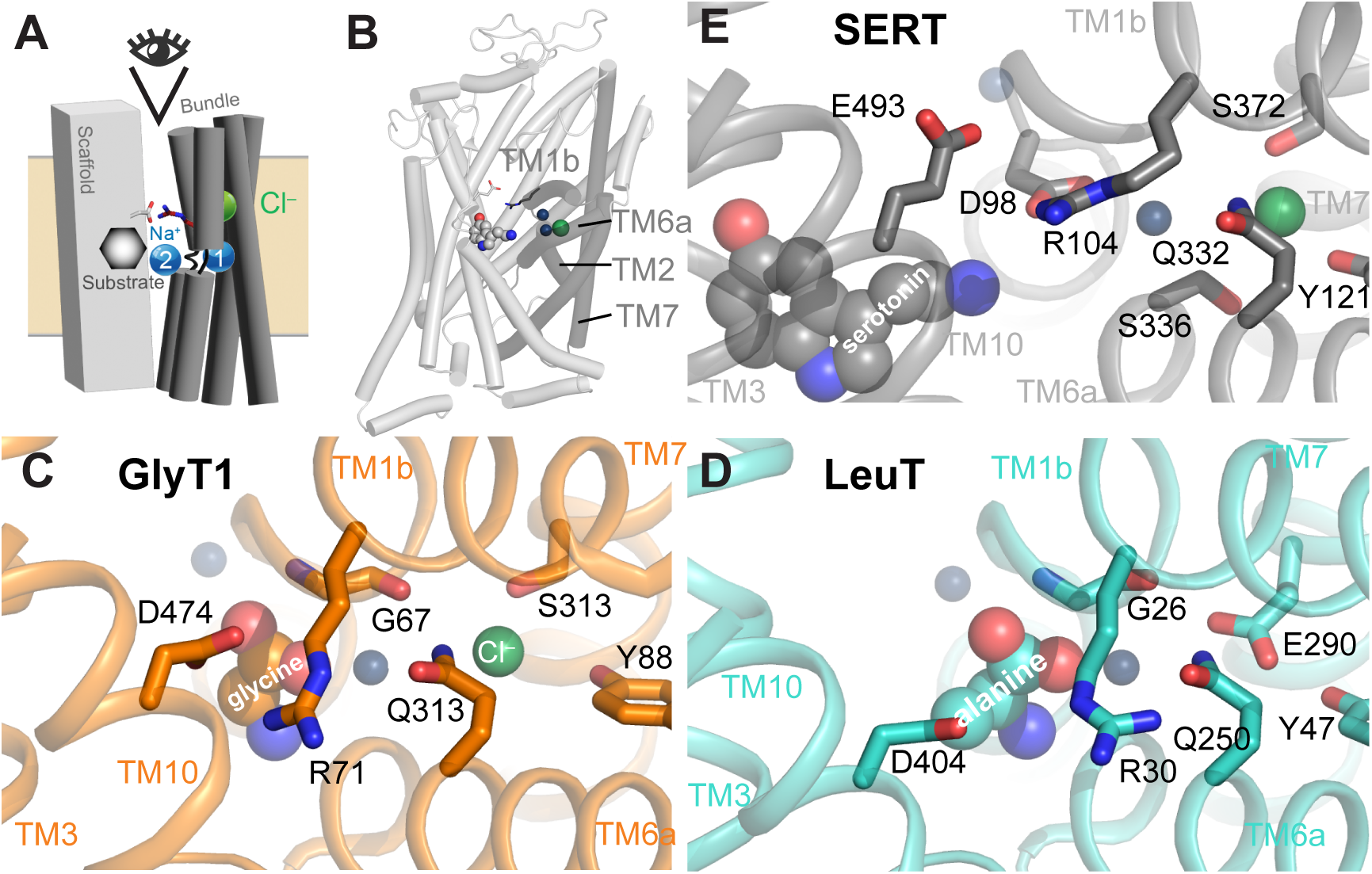
SLC6 transporter structures. **(A, B)** Schematic and cartoon representation of the LeuT fold, illustrating the central locations of the substrate (*hexagon, gray spheres*), Na^+^ (*blue spheres*) and Cl^−^ ions (*green spheres*), and the position of the extracellular gate salt bridge residues (*sticks*). The bundle region, comprising transmembrane helices 1, 2, 6 and 7 is in dark gray, while the scaffold region is in light gray. Oxygen atoms are colored red, nitrogen in blue. **(C-E)** The extracellular interaction network viewed from the orientation indicated in panel A. The Gln-Arg-Asp/Glu network connecting the Cl^−^ binding site to helices in the scaffold is highlighted in sticks, as are Cl^−^ binding residues. In the amino acid transporters GlyT1 (C; PDB entry 8WFI) and LeuT (D; PDB entry 3F48), the key salt bridging acidic residue in TM10 is an aspartate (Asp474 and Asp404, respectively), whereas in the monoamine transporter SERT (E; PDB entry 7LIA), Glu493 plays this role. In LeuT, the Cl^−^ ion is replaced by the side chain of Glu290. Protein is shown as cartoon helices viewed from the extracellular side. Ions and substrates are shown as spheres, and key residues are shown in stick representation and labelled using the numbering of the corresponding transporter. Note that the sequence numbering in GlyT1b here is that in the structure [21], UniProt identifier P48067-3, a different isoform from that referred to in the main text [20]. Amino acid substrates provide a carboxylate group to interact with a tyrosine in TM3 and the Na^+^ ion at the Na1 site, whereas the monoamine substrates of SERT, NET and DAT lack this carboxylate group, which is replaced by the γ-carboxylate of an aspartate side chain in TM1 (Asp98 in SERT). Structures are rendered using PyMol (Schrödinger, Inc).

Early results demonstrated that Cl^−^ was required for 5-HT uptake into platelets (13), plasma membrane vesicles from platelets (14), synaptosomes (15), brush-border membrane vesicles (16), and cells transfected with SERT cDNA (17). Although these results were consistent with stoichiometric Cl^−^ symport with 5-HT^+^ (18), more recent data indicates that intracellular Cl^−^ does not affect transient currents associated with 5-HT^+^ transport, suggesting that intracellular release of chloride may not be coupled to that of 5-HT^+^ and Na^+^ (19). Consequently, SERT might diverge from the NSS amino acid transporters for glycine (GlyT1) and GABA (GAT-1), which couple substrate and Cl^−^ influx (20, 21). Nevertheless, Cl^−^ is required for SERT function (13), raising the question of how Cl^−^ mediates its influence on SERT. Cool et al reported that Cl^-^increased the affinity of 5-HT, measured as the ability to displace bound β-CIT (16). An additional likely contribution is that Cl^−^ binding increases the affinity of SERT for Na^+^ (22). Previous studies of GlyT1 revealed another influence of Cl^−^, namely on the conformational state of the transporter (23). Specifically, in the presence of Na^+^, addition of either the substrate (glycine) or Cl^−^ alone led to a decrease in accessibility in the extracellular permeation pathway. However, these combinations of substrates resulted in much smaller changes in the cytoplasmic pathway, which is stabilized in a closed conformation by Na^+^. When both glycine and Cl^−^ were added together with Na^+^, there was a further decrease in accessibility of the extracellular pathway and a large increase in the cytoplasmic pathway, consistent with the conversion from outward- to inward-facing conformation (24). These results also suggested that substrate and Cl^−^act independently to affect conformational change in GlyT1.

The accessibility data for GlyT1 suggest that Cl^−^ may influence the conformational state also for SERT. However, despite the similarities in sequence and structure between SERT and GlyT1, there are important mechanistic differences. For example, GlyT1 transports two Na^+^ ions and a Cl^−^ ion together in the step that delivers glycine to the cytoplasm (20), whereas there is evidence that SERT transports only one Na^+^ with 5-HT^+^ (6). Because Cl⁻ lies adjacent to the Na1 site, differences in Na^+^ handling between NSS transporters could have direct consequences on the behavior of Cl⁻ during the cycle. SERT also utilizes the outwardly directed K^+^ gradient to drive the return step (7, 8) while GlyT1 apparently does not couple K^+^ and glycine fluxes (25). Substrate interactions also differ markedly. For amino acid transporters in the NSS family, such as GlyT1, an important interaction (first discovered in a bacterial amino acid transporter, LeuT) is formed between the substrate carboxyl group and one of the two bound Na^+^ ions (**Fig. 1C, 1D**) (24, 26, 27). However, amine substrates of transporters like SERT, such as 5-HT^+^, lack this carboxyl and in NSS amine transporters like SERT, a protein aspartate side chain γ-carboxyl (as in Asp98 in SERT) occupies a similar position to that of amino acid substrates (**Fig. 1E**) (27, 28). Other NSS amine transporters (for norepinephrine and dopamine; NET and DAT, respectively) also contain aspartate at the position corresponding to Asp98 in SERT (29–32), whereas all NSS amino acid transporters contain glycine at that position (**Fig. 1C**).

Given the known mechanistic differences between amino acid and monoamine transporters, we considered whether Cl^−^ acts similarly in GlyT1 and SERT. From a structural perspective, the interactions in the Cl^−^ binding site in SLC6 transporters are well conserved, as is the interaction network connecting it to the extracellular pathway (**Fig. 1C-E**). For example, a conserved glutamine residue (Gln332 in SERT) interacts with the bound Cl^−^ ion in both transporters (23, 33). In chloride-independent LeuT (and in GlyT1), this glutamine is close to a conserved arginine (Arg104 in SERT), which alternatively can salt bridge with an acidic residue across the extracellular pathway (Glu493 in SERT) (26, 28), depending on the orientation of the arginine side chain (**Fig. 1C-E**). Formation of this salt bridge is thought to contribute to closure of the extracellular pathway in NSS transporters (34, 35) including SERT (36). Thus, we proposed for GlyT1 that the effect of Cl^−^ binding shifted the arginine rotamer distribution toward formation of this salt bridge by favoring interaction of the arginine with the aspartate, rather than the glutamine (see **Fig. 1C**). Mutation of this glutamine to glutamate stabilized GlyT1 in an outward-open state (as measured by decreased accessibility of the cytoplasmic pathway) and was consistent with a stronger interaction between the arginine and glutamate, relative to the wild-type glutamine that it replaced. (Note that due to differences in sequencing and isoforms, Gln313 in Figure 1C corresponds to Gln299 in the work of Zhang et al. (23).)

Here, we revisit the role of Cl^−^ in the mechanism of SERT. First, to assess whether Cl^−^ and substrate movements are coupled, we measure uptake into SERT proteoliposomes in the presence and absence of a Cl^−^ gradient. Second, using accessibility measurements and molecular dynamics simulations, we examined whether Cl^−^ influences conformation in SERT in the same way as in GlyT1. The findings explain why Cl^−^ is required for transport and demonstrate which features unique to the sequence of SERT are important in mediating chloride’s effect.

## Results

### Effect of transmembrane Cl^−^ concentration gradients on SERT-mediated substrate influx

In transporters, coupling requires that a gradient of one transported species be capable of driving the net movement of another. Although Cl^−^ is required for SERT-mediated 5-HT^+^ transport, previous studies have not conclusively demonstrated that Cl^−^ gradients can provide a driving force for substrate accumulation. Instead, coupling was inferred from (i) the requirement for external (but not internal) Cl^−^ (18) and (ii) evidence that the 5-HT^+^ transport cycle by SERT was electrically neutral (7). Subsequent electrophysiological studies showing that intracellular Cl^−^does not influence transport-associated currents called into question whether Cl^−^ is co-transported with 5-HT^+^ (19). To directly assess whether Cl^−^ gradients can drive 5-HT^+^ uptake, we reconstituted purified SERT into proteoliposomes (9), allowing direct measurements of substrate accumulation in the presence and absence of Cl^−^ gradients.

We generated SERT proteoliposomes using the detergent-free DIBMA method (9, 37). Proteoliposomes prepared in one medium were diluted into another medium containing [^3^H]5-HT, allowing independent control of imposed transmembrane ion gradients. Except where indicated, solutions were matched for osmolarity and ionic strength and differed only in the identity and distribution of the ions used to establish defined gradients. When SERT proteoliposomes generated in high-K^+^ were diluted into high-Na^+^ medium, accumulation of 5-HT^+^ followed a rapid rise driven by the imposed Na^+^ and K^+^ gradients, reaching maximal levels at 1.5-5 mins, followed by a slow (t_1/2_ 15-20 min) decay characteristic of dissipation of the ion gradients with time (**Fig. 2A**, black circles). To assess any nonspecific association of substrate to the liposome surface, we applied gramicidin, which increases membrane permeability to K^+^ and Na^+^ leading to collapse of these ion gradients. When added to the K^+^-loaded proteoliposomes 5 min. prior to dilution into Na^+^ medium with 5-HT^+^, gramicidin almost completely ablated 5-HT^+^ accumulation (**Fig. 2A**, filled red circles). When gramicidin was added to the dilution medium (containing Na^+^ and 5-HT^+^), accumulation was markedly decreased, and the rate of decay markedly accelerated (**Fig. 2A**, open red circles), reinforcing the conclusion that 5-HT^+^ accumulation was driven by the imposed ion gradients and not an artifact of 5-HT binding. The increased effectiveness of gramicidin when pre-incubated with the proteoliposomes likely resulted from its higher relative concentration and the time needed for formation of the conductive gramicidin dimer during the 5 min. pre-incubation (38); in either case, the results are consistent with gramicidin’s action on Na^+^ and K^+^ permeability and the known influence of those gradients on 5-HT^+^ transport.

**Fig. 2.**
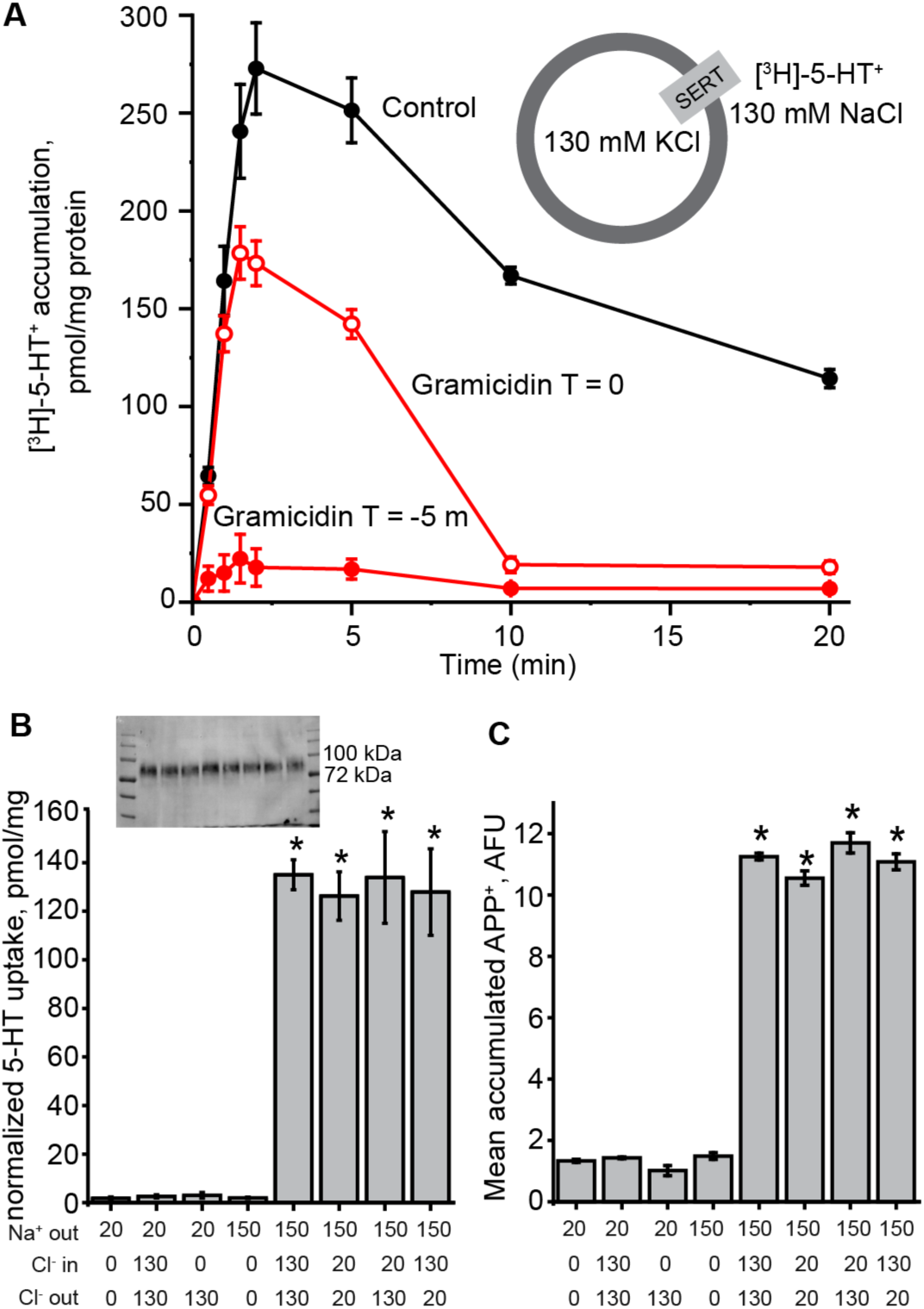
Influence of Na^+^ and Cl^−^ gradients on substrate accumulation by reconstituted SERT liposomes. (**A**) Time course of [^3^H]-5-HT^+^ accumulation into SERT liposomes in the presence of an inward-directed Na^+^ gradient. Reconstituted SERT proteoliposomes (200 ng protein) in 130 mM KCl containing 10 mM K_2_HPO_4_ were diluted 20-fold into 190 µl of 130 mM NaCl containing 10 mM Na_2_HPO_4_ and 20 nM [^3^H]5-HT as shown in the inset, both solutions adjusted to pH 7.4 with phosphoric acid. Gramicidin, where added, was present at a final concentration of 50 µM in the assay mixture. Black circles, no gramicidin; Red symbols, gramicidin present at 50 µM in the reaction mixture; Open red circles, gramicidin added to external (Na^+^) solution before addition of proteoliposomes; Filled red circles, gramicidin added to proteoliposome suspension (K^+^) 5 min before dilution. Accumulated 5-HT was measured as described in “Materials and Methods”. (**B**) Effect of imposed ion gradients on peak 5-HT accumulation. SERT proteoliposomes were formed in 10 mM Na_2_HPO_4_, (pH 7.4) with 130 mM of either the gluconate or Cl^−^ salt of NMDG^+^ (N-methyl-D-glucamine), , and diluted into 10 mM Na_2_HPO_4_, (pH 7.4) with 130 mM either Na^+^ or NMDG^+^ salt of either or both Cl^-^ or gluconate to give the final concentrations indicated in the x-axis legend (see **Table S3**) and to maintain the same ionic molarity while generating inward-directed gradients of Na^+^ and Cl^−^. The dilution solution also contained 20 nM [^3^H]5-HT. After 2 min, 5-HT accumulation was determined as above. Accumulated [^3^H]5-HT^+^ was normalized to the level of SERT determined using Western Blot (inset) for each condition. Western blot intensities were all within 15% of control (first lane), averaging 88.1±11.6% over 3 separate experiments. Asterisks indicate significantly different levels of 5-HT^+^ accumulation, relative to those measured in the absence of Na^+^ gradient or at 0 Cl^−^ (columns 1-4) (*P < 0.05) using Student’s paired t test. All error bars represent the SEM; n = 3. None of the [^3^H] 5-HT^+^ levels in columns 1-4 were significantly different from each other (P > 0.44). Differences between columns 5-8 were also not significant (P > 0.2). (**C**) Fluorescence levels measured for APP^+^ substrate uptake under the same conditions for [^3^H]5-HT uptake. Asterisks and error bars have the same meanings as in panel B. None of the APP^+^ levels in columns 1-4 were significantly different from each other (P > 0.35). Differences between column 5 *vs.* 6, 7 or 8 were not significant (P > 0.44), and P values for the differences between columns 6 *vs.* 7, 6 *vs.* 8 and 7 *vs.* 8 were between 0.076 and 0.125.

The ability of a Cl^−^ concentration gradient to drive substrate accumulation was tested by varying the ion concentrations and normalizing the level of 5-HT^+^ obtained at 2 mins to the amount of protein detected on a Western blot (**Fig. 2B**). The 2 min time-point was chosen as it precedes substantial dissipation of Na^+^/K^+^-driven accumulation. In the absence of a transmembrane Na^+^ gradient, with ionic strength maintained by NMDG^+^ (leftmost 3 columns; Na^+^ was 20 mM in every solution), no significant accumulation of the 5-HT^+^ occurred when Cl^−^ was absent (column 1), present on both sides at equal concentrations (column 2), or present only outside the vesicles (column 3). The presence of a Na^+^ gradient (columns 4-8) alone was not sufficient to increase accumulation in the absence of Cl^−^ (column 4), but when Cl^−^ was present outside the vesicle, accumulation increased. This increase was similar in the presence of either 20 or 130 mM external Cl^−^ but was independent of the imposition (compare columns 5 or 6 with 7 or 8) or direction (columns 7 vs 8) of a transmembrane Cl^−^ gradient. Because ion gradients were imposed under otherwise identical conditions, any effects of Cl⁻ on accumulation would reflect coupling to transport rather than indirect effects on membrane potential.

The relative amounts of SERT in the proteoliposomes was determined for this experiment (shown in inset of **Fig. 2B**). Although only correctly-oriented and active transporters contribute to uptake, the absence of condition-dependent differences in SERT incorporation indicates that differences in accumulation are not attributable to altered protein reconstitution. These findings with 5-HT^+^ were recapitulated by measuring accumulated fluorescence of the commonly used substrate analog APP^+^ (**Fig. 2C**), indicative that both substrates were accumulated in response to the same transmembrane ionic gradients.

To control for the unlikely possibility that the proteoliposomes used in these experiments were so leaky that the imposed Cl^−^ gradients dissipated before 5-HT^+^ accumulation could occur, we tested the ability of Cl^−^ to dissipate a membrane potential generated by valinomycin (a K^+^-specific ionophore) combined with an imposed K^+^ gradient. Proteoliposomes prepared in high KCl or K-gluconate were diluted into isosmotic Na-gluconate after which di-S-C3-(5), a permeant fluorescent cation, was added to a final concentration of 1 µM. Di-S-C3-(5) accumulates inside vesicles or cells when the interior is electrically negative relative to the external solution (39), and this accumulation quenches its fluorescence. Upon addition of valinomycin to liposomes with a transmembrane K^+^ difference (high in, low out), K^+^ efflux occurs until a membrane potential (negative inside) is established that counteracts further K^+^ efflux. Compensating ion fluxes, such as Cl^−^ efflux, can collapse the potential (coupled KCl efflux), leading to recovery of fluorescence.

The establishment of such a K^+^ diffusion potential in liposomes without SERT or DIBMA is shown in **Fig. S1** (lower traces). diS-C3-(5) added at time 0 created a baseline fluorescence that was decreased when valinomycin was added at 2 min, indicating generation of a membrane potential (negative inside). Detergent (Triton x-100, 1% final) was added at 8 min, to disrupt the vesicles and release the dye, increasing its fluorescence. Notably, the fluorescence traces were identical in the presence or absence of intraliposomal Cl^−^ (red vs. black symbols). Similar results were obtained using proteoliposomes reconstituted with SERT-DIBMA (middle trace), and again internal Cl^−^ did not lead to potential collapse relative to gluconate, an impermeant anion. The upper trace shows that fluorescence in SERT proteoliposomes was insensitive to valinomycin if no K^+^ gradient was generated. These results are consistent with the observation that here, and in previous work, Na^+^, K^+^, and H^+^ gradients were preserved long enough to support measurable accumulation of 5-HT or APP^+^ (**Fig. 2A** and [Ref. (9)]). For significant net Cl^−^ flux to occur, compensating flux of one of these other ions would need to occur. Together the results argue strongly that Cl^−^ permeability in these SERT proteoliposomes was low, and certainly no greater than that of gluconate.

Because Cl⁻ is required for accumulation but does not provide a driving force, an alternative possibility is that Cl⁻ influences transport by stabilizing ion binding or conformational equilibria. Consistent with the former idea, the Cl^−^ ion is only ∼8 Å from the Na1 site cation in NSS transporter structures, and Na^+^ and Cl^−^ affinities are mutually dependent (22). In line with a role in conformational equilibria, Cl^−^ impacts binding of the SERT inhibitor fluoxetine (Prozac)(40). However, the conformational effects of chloride of SERT had not been measured directly.

### Conformational effects of Cl^−^ and 5-HT^+^ on SERT

To assess conformational shifts in SERT, we measured the reactivity of cysteine residues replacing either Ser277 in the cytoplasmic pathway or Tyr107 in the extracellular pathway (**Fig. 3, Fig. S2, Table S1**). Changes in cysteine reactivity at these positions report on the openness of the cytoplasmic and extracellular permeation pathways, respectively, which have been shown to correlate with inward- and outward-facing conformational states in LeuT, GlyT1, and SERT (10, 23, 27). As seen in outward- and inward-facing SERT structures, Tyr107 was more accessible to the extracellular medium in outward-facing SERT conformations (**Fig. 3E**) and Ser277 was more accessible to the cytoplasm in inward-facing conformations (**Fig. 3F**). Reactivity of cysteine residues replacing Ser277 or Tyr107 in SERT or the corresponding residues in LeuT and GlyT1, have been successfully used to access accessibility of the extracellular or cytoplasmic permeation pathway, respectively (5, 10, 23, 41–47).

**Fig. 3.**
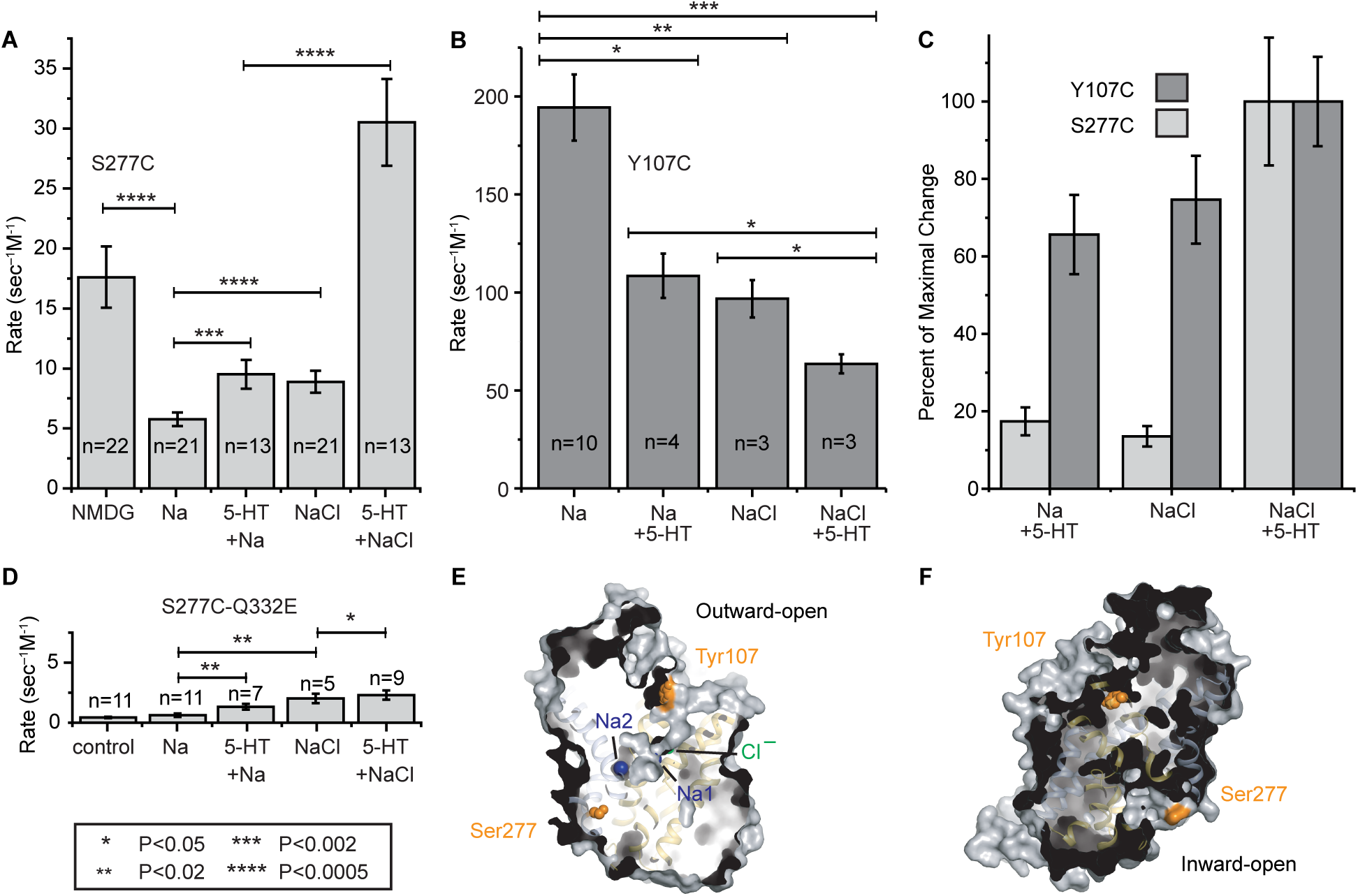
Accessibility changes in SERT induced by ions and substrate serotonin. (**A, B**) The conformational response to ligand binding was estimated by changes in the reactivity of Y107C in the extracellular pathway and S277C in the cytoplasmic pathway toward MTSEA added to the medium of intact cells expressing SERT Y107C-C109A (74) or to membrane samples from cells expressing SERT S277C-X5C (10). Unless explicitly labelled “Cl”, Cl^−^ was replaced by gluconate. Concentrations were 0.1 M for Na^+^ or NMDG^+^ and 5-HT was added at 10 µM. (**C**) Conformational response, in the presence of Na^+^, to Cl^−^ or 5-HT^+^, shown as relative to the maximal response in each case. The change in accessibility of Y107C and S277C between Na^+^ alone and NaCl with 5-HT^+^ was set to 100% and the changes induced by Cl^−^ or 5-HT^+^ scaled accordingly. Light grey bars represent cytoplasmic pathway (S277C) reactivity and dark grey bars, that of the extracellular pathway (Y107C). All error bars represent the SEM; n ≥ 4. Note that some of these differences may arise from the different conditions required to assay the two pathways. **(D)** S277C reactivity with additional mutant Q332E. Asterisks indicate significantly different rate constants between columns at the ends of the horizontal bars (*P < 0.05; **P < 0.02, etc. as indicated) using Student’s t test. Error bars represent the SEM; n ≥ 4. (**E, F**) Surface exposure of residue Tyr107 and Ser277 (*orange spheres*) in SERT shown for outward-facing (A) and inward-facing (B) conformations, PDB entries 7LIA and 6DZZ, respectively, viewed from within the membrane plane. The protein is shown as surface and helices, with the bundle (TM1, 2, 6, 7) in yellow, and other helices in gray; Na^+^ (*blue*) and Cl^−^ions (*green*) are shown as spheres where present.

In the cytoplasmic pathway, Cys277 accessibility was measured in membrane fragments, which allow access of the sulfhydryl reagent MTSEA (2-Aminoethyl methanethiosulfonate) to the cytoplasmic side. The resultant cysteine modification inhibited ligand binding, measured here with [^125^I] β-CIT. In these measurements and in intact cells (below), the MTSEA concentration at 50% inhibition and the reaction time were used to calculate pseudo first-order rate constants. Under these conditions, decreased reactivity indicates stabilization of the cytoplasmic pathway in a closed state. SERT-S277C reactivity was decreased in the presence of Na^+^ (**Fig. 3A**, column 1-2) as previously reported (2), reflecting Na^+^-dependent stabilization of the closed cytoplasmic pathway in SERT and other NSS transporters (3, 10, 23, 48, 49). In the presence of Na^+^, addition of either 5-HT^+^ (column 3) or Cl^−^ (column 4) partially reversed the effect of Na^+^, and addition of both 5-HT^+^ and Cl^−^ further increased cytoplasmic pathway accessibility to an extent greater than in the absence of Na^+^ (column 5).

In the extracellular pathway, Cys107 accessibility was measured in intact cells using the effect of MTSEA on [^3^H]5-HT transport activity after washing. The response to 5-HT^+^ and Cl^−^ was opposite to that observed for the cytoplasmic pathway (**Fig. 3B**). Specifically, in the presence of Na^+^ (column 1), addition of 5-HT^+^ markedly decreased accessibility of Cys107 (column 2) and addition of Cl^−^ produced a similar effect (column 3). Addition of both 5-HT^+^ and Cl^−^ further decreased accessibility (column 4).

Although cytoplasmic and extracellular accessibility were measured using distinct assay formats, normalization to the maximum effect observed with both ligands allows comparison of how 5HT^+^ and Cl^−^ bias modulation of each pathway (**Fig. 3C**). Addition of 5-HT^+^ or Cl^−^ changed cytoplasmic accessibility (light gray bars in the first two pairs of columns) by only about 15-20% of the maximal effect, whereas the same ligands altered accessibility of the extracellular cysteine by 65-70% of maximal change (dark gray bars in the first two pairs of columns). The distinct assay formats for measuring extracellular and cytoplasmic accessibility require a caveat that intact cells maintain transmembrane gradients of Na^+^, Cl^-^ and K^+^, while the membrane fragments do not. It is possible that increased Na^+^ and 5-HT on the cytoplasmic face of the membrane fragments could bias the conformational equilibrium toward outward-facing states, but the robust response of SERT to Cl^-^ and 5-HT suggests that these influences do not prevent the ability of Cl^-^and 5-HT to open the cytoplasmic pathway.

### Behavior of an extracellular pathway network depends on the state of the transporter

To identify the molecular interactions responsible for measured conformational dependence, we turned to Molecular Dynamics (MD) simulations. In our previous study using LeuT (23), we found that two conserved residues, Arg30 and Gln250 (corresponding to Arg104 and Gln332 in SERT, respectively, **Fig. 1**) interacted consistently in the outward-open state of LeuT, but not in the outward-occluded state, where Arg30 instead salt-bridges to Asp404 (corresponding to Glu493 in SERT), forming the so-called extracellular gate (**Fig. 1D**). Thus, the Arg30-Asp404 salt bridge appears to be a feature of, or to influence the degree of, extracellular closure (and consequently also of intracellular opening). Note that in those simulations, the nearby Glu290 in LeuT was in its charged state to mimic the Cl^−^-bound network of SERT. Based on those observations, it was proposed that Cl^−^ influences salt-bridge formation and pathway closure, and that it does so via the conserved Gln residue (3, 23, 50).

Here, we carried out simulations of SERT with bound 5-HT^+^, Na^+^, and Cl^−^ and computed the distances and H-bond formation between Arg104 and Gln332 or Glu493. Simulations were initiated from defined outward-open or outward-occluded states and were designed to sample these states rather than interconversion between conformations. Consistent with the findings for LeuT, Arg104 of SERT could H-bond with Gln332 in the outward-open state (**Fig. 4A-B; Fig. S3A**) but not in the outward-occluded state (**Fig. 4C-D; Fig. S3C**). Indeed, in the outward-open state, Arg104 was within H-bonding distance of Gln332 for around 60% of the aggregated simulation time (**Fig. 4A**), reflecting a preferred interaction in this conformational state, albeit with variability between trajectories (**Fig. S3A**). This variability reflects substantial dynamics of these helices and side chains in the outward-open state, which are apparent upon visualizing snapshots from each trajectory (**Fig. 5**).

**Fig. 4.**
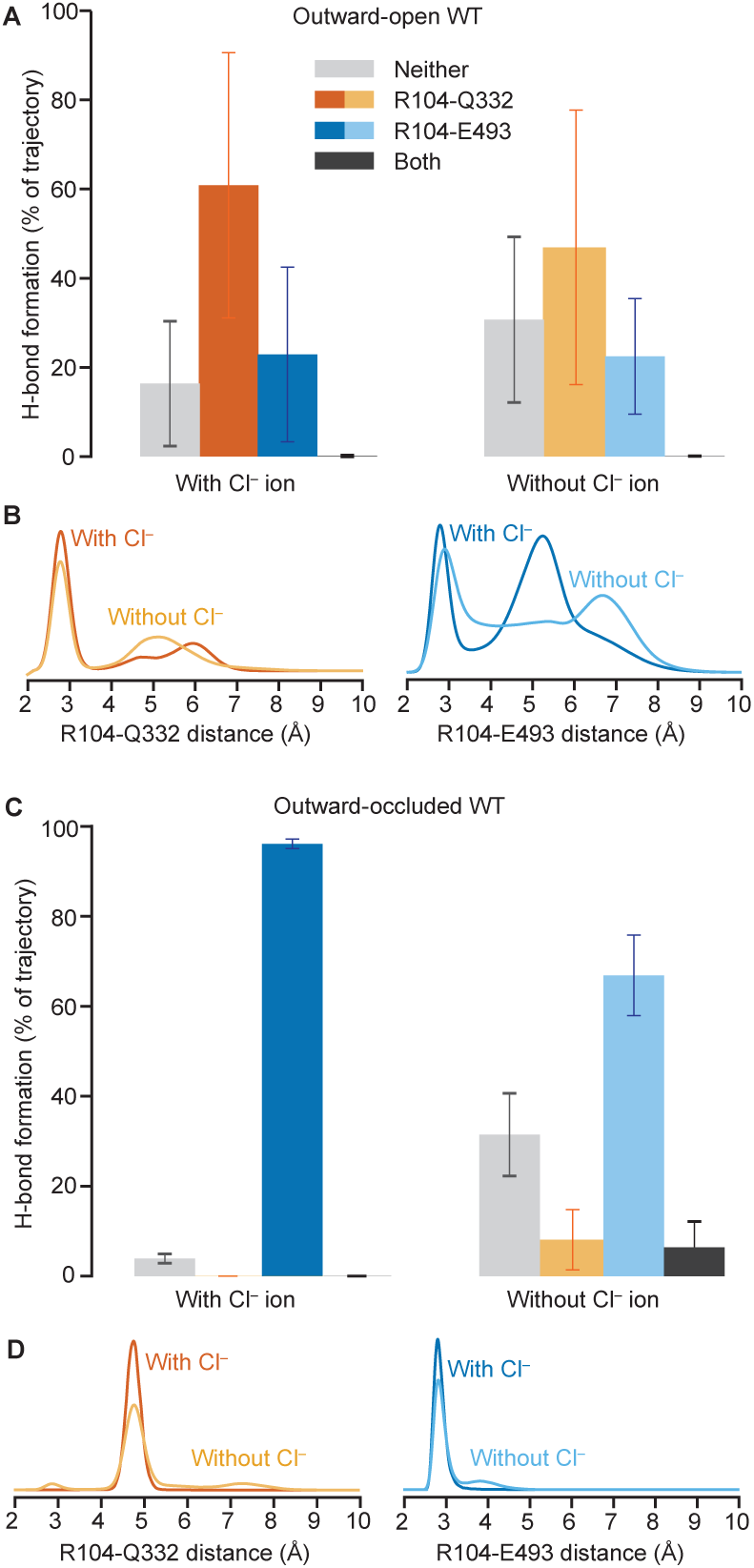
Behavior of the interaction network connecting the Cl^−^ binding site to the extracellular pathway salt bridge in wild type SERT. Substrate 5-HT^+^ and two Na^+^ ions were placed within their respective sites. Molecular dynamics simulations were initiated with **(A, B)** outward-open and **(C, D)** outward-occluded conformations for 0.5 µs simulations (n=4 or n=8, respectively). **(A, C)** Bar charts indicate the fraction of the aggregate simulation time that a H-bond or salt bridge (defined as distance <3.2 Å) can be formed. The atom selections included any donor atom of Arg104 and any acceptor atom of either Gln332 (*orange bars*), Glu493 (*blue bars*) or both (*black bars*), either in the presence (*left*) or absence (*right*) of a Cl^−^ ion initially placed at its reported binding site. Gray bars represent the fraction of time that neither interaction is formed. Error bars reflect standard deviations across repeat trajectories. The large error bars in panel A compared to C reflect variability between trajectories of the outward-open state where the dynamics are more substantial. (**B, D**) We combine the data over all trajectories into a distribution of minimum distances for SERT simulations either in the presence (*darker lines*) or the absence (*lighter lines*) of the Cl^−^ ion and the differences between states reflect individual trajectory data, shown in **Fig. S3**.

**Fig. 5.**
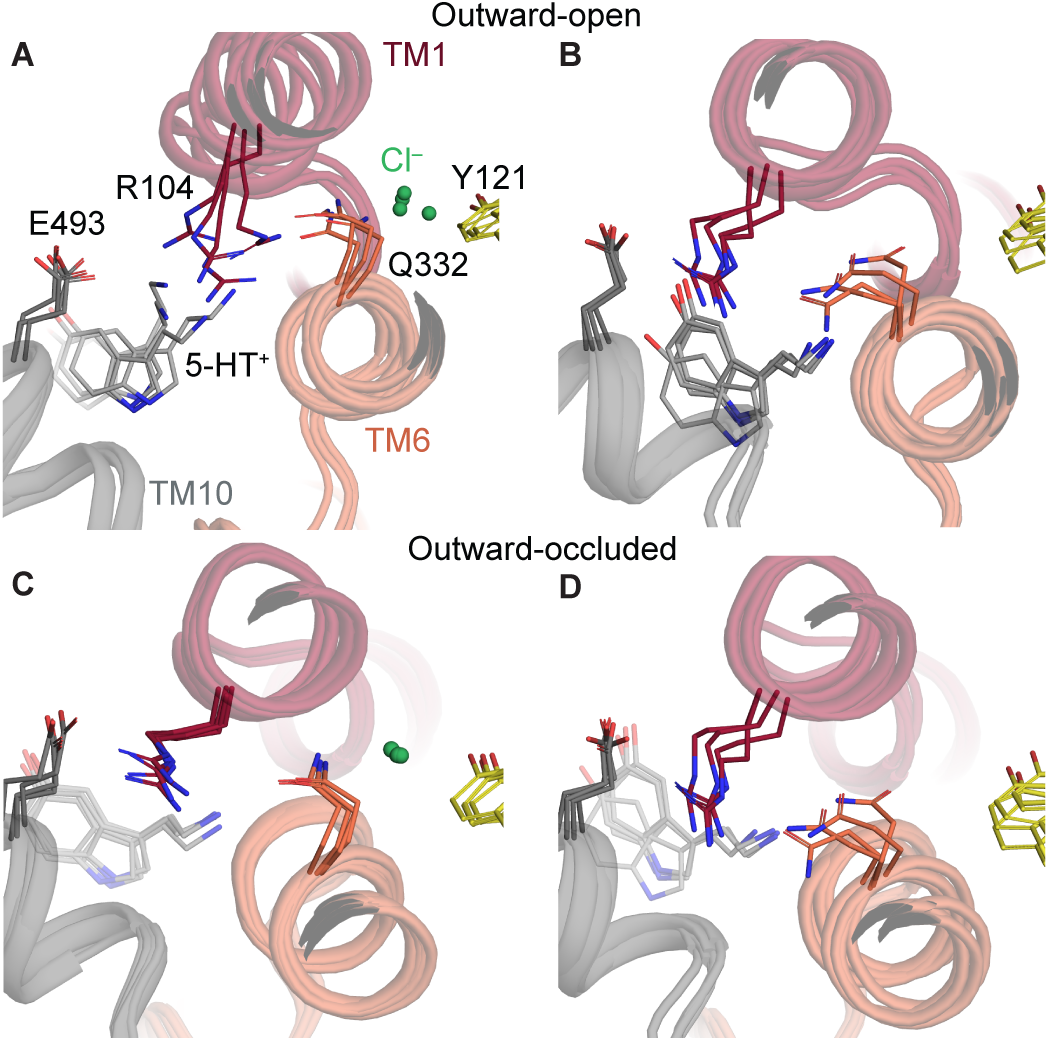
Interaction network connecting the extracellular pathway with the Cl^−^ site in SERT. Simulation snapshots are viewed from the extracellular side, as indicated in Figure 1A. (**A-D**) Multiple frames were taken from simulations of the **(A, B)** outward-open and **(C, D)** outward-occluded conformations, either in the presence (A, C) or absence (B, D) of a Cl^−^ ion (*green sphere*), with TM1 (*red*), TM6 (*orange*) and TM10 (*gray*) shown in cartoon representation. Key residues and the substrate 5-HT^+^ (*grey*) are shown as sticks with O atoms in red and N atoms in blue. Tyr121 from TM2 is colored yellow.

Further consistency with LeuT is seen in that Arg104 of SERT formed a salt bridge with the acidic residue Glu493 more frequently in the outward-occluded state (**Fig. 4C-D; Fig. S3G**) than in the outward-open state (**Fig. 4A-B; Fig. S3E**). These simulations of Cl^−^-bound SERT in different conformations are consistent with the possibility that Gln332 couples Cl^−^ binding to changes in the interactions that govern closure of the extracellular pathway salt-bridge.

### Behavior of the extracellular pathway in the absence of a Cl^−^ ion

In LeuT, the absence of chloride was, in one study, mimicked by protonation of Glu290 (50), which is an imperfect analog. To distinguish whether the observed interaction network is a consequence of Cl^−^ binding itself, or the underlying protein architecture, we repeated the simulations of SERT without the bound Cl^−^ ion.

In simulations of SERT with the extracellular pathway open, the Gln332-Arg104-Glu493 network behaved similarly whether Cl^−^ was bound or not (**Fig. 4A**, left vs right; **Fig. 4B**, dark vs light colors), even when accounting for the heightened dynamics of this state (**Fig. S3A-F**). For example, the Arg104-Glu493 salt bridge interaction (**Fig. 4A-B**, blue) was formed around 20% of the aggregate simulation time both with and without the Cl^−^ ion. Thus, the anion did not substantially modify the local network for the outward-open state.

By contrast, in simulations with the extracellular pathway closed, the local network was influenced by the ion (**Fig. 4C-D**). In particular, the side chain of Gln332 was more dynamic in the absence of the Cl^−^ ion (**Fig. S3D; Fig. 5D**) than when the ion was bound (**Fig. S3C; Fig. 5C**). To describe the internal rotations of the Gln332 side chain, we measured the χ_1_ dihedral angle; this angle remains in *trans* conformation in the presence of the Cl^−^ ion but rotates from *trans* toward *gauche*(+) in the absence of the ion in multiple trajectories (**Fig. S4B**). In the latter orientation, Gln332 can interact with Arg104 (**Fig. 5D; Fig. S3D**), which would be expected to favor loss of the Arg104-Glu493 salt bridge (**Fig. 5D; Fig. S3H**). However, Arg104 remains within 4 Å of Glu493, even when the Cl^−^ ion is absent (**Fig. S3G-H**). While longer than the typical distance of a salt bridge, the two charges would still experience each other at this distance in these atomistic simulations. Accordingly, despite stabilizing the local network in the occluded state more than in the outward-open state (and therefore being consistent with a role in favoring occlusion), the anion does not act exclusively via Gln332-mediated enhancement of salt-bridge formation in SERT, as previously proposed.

### Chloride acts on features other than the extracellular salt bridge in SERT

To identify potential broader consequences of the absence of a Cl^−^ ion that could further destabilize the occluded state and lead to the measured accessibility changes, we characterized the dynamics of nearby residues. The most notable changes in dynamics were observed for Asp98, whose side chain χ_2_ angle predominantly adopted the *trans* orientation seen **Fig. 1E** in the simulations with Cl^−^ bound (**Fig. S4A**). When Cl^−^ was absent, however, the Asp98 side chain could rotate freely (**Fig. S4B**), despite being within salt bridge distance of the charged primary amine of the bound 5-HT^+^ ion and, on the other side, to the Na^+^ ion in the Na1 site, which might be expected to stabilize the carboxyl position.

The increased dynamics of Asp98 and other regions relative to the Cl^−^-bound protein coincide with greater hydration of this region (**Fig. 6**). Specifically, water molecules from the extracellular side solvate not only the empty Cl^−^ site, but also Asp98, the extracellular salt bridge region of the pathway, and other regions within the bundle helices (**Fig. 6B**). We therefore posit that increased conformational freedom – as exemplified in **Fig. 5** – contributes to reduced pathway closure in the absence of Cl^−^ ions.

**Fig. 6.**
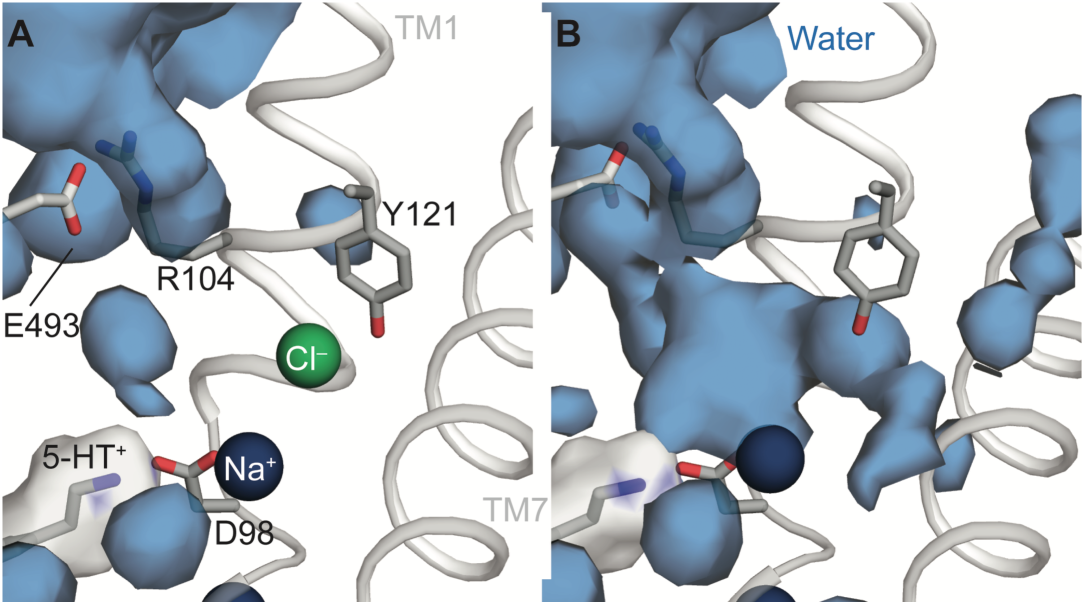
Hydration of the outward-occluded conformation of SERT increases in the absence of a Cl^−^ ion. Water occupancy maps (*blue surface*) are shown with an equilibrated conformation of SERT as reference, for Molecular Dynamics simulations of the outward-occluded state in **(A)** the presence and **(B)** the absence of a Cl^−^ ion (*green sphere*). Occupancies within 20 Å of Asp98 were averaged over n = 8 trajectories, respectively. Selected side chains and 5-HT^+^ are shown as sticks with O atoms colored red and N atoms in blue. The Na^+^ ion is in the Na1 site (*blue sphere*) and the occupancy map of 5-HT^+^ is shown (*white surface*). Tyr121 is from TM2, whose helical backbone is omitted for clarity.

To assess whether the propensity for extracellular pathway closure is reduced in the absence of Cl^−^, we computed the distance between helices in the bundle (either TM1b or TM6a) and a reference helix in the scaffold, namely TM9 (**Fig. S5**). For the outward-open conformation of SERT with Cl^−^ bound, the initial ∼28 Å separation between TM1b and TM9 decreases over time (**Fig. S5A**) and starts to resemble outward-occluded structures (∼26.5 Å). The TM6a-TM9 distance remains around 29 Å (**Fig. S5B**), consistent with a previous simulation study that found TM1b closes prior to TM6a (51). However, in the absence of the ion both TM1b and TM6a tended to remain at their initial positions, with fewer transitions to short distances consistent with closure (**Fig. S5A-B**). Conversely, there were signs that TM6a starts moving away from TM9 in the simulations of the occluded state without the anion, indicative of pathway opening (**Fig. S5D**).

These observations provide a structural explanation for the increased extracellular accessibility observed experimentally (**Fig. 3**): for wild-type SERT in the presence of Na^+^ alone, or both Na^+^ and 5-HT^+^, the addition of Cl^−^ reduced the accessibility of Y107C in the extracellular pathway consistent with reduced hydration and closer helix packing in the presence of Cl⁻. These effects are likely to be cumulative with the reduced Na^+^ binding affinity measured in low Cl^−^concentrations implied by the data in Tavoulari et al (22). We conclude that the three distinct effects: (i) reduced sodium affinity; (ii) reduced salt-bridge formation; and (iii) reduced stability of nearby sidechains, together significantly diminish the probability of forming the occluded conformation in the absence of chloride, likely increasing the kinetic barrier to transport.

### Testing the proposed salt-bridge stabilization by chloride

The contribution of the extracellular salt bridge to the chloride response was previously tested in GlyT1, by modifying the glutamine that links chloride binding to the salt-bridge arginine. Specifically, Gln299 (corresponding to SERT Gln332, **Fig. 1**) was changed to glutamate, as it was proposed that a negatively charged carboxyl side chain at this position would sequester Arg57 (Arg104 in SERT) and prevent it from establishing the salt bridge with Asp460 (Glu493 in SERT). Indeed, in GlyT1, this mutation led to a dramatic decrease in the accessibility of residues in the cytoplasmic pathway, as if the transporter were strongly biased toward inward-closed conformations. We tested the corresponding mutation in SERT, with similar overall results (**Fig. 3D**) (note the similar y-axis scales in **Figs. 3A** and **3D**). As in GlyT1, this mutation and the resulting conformational effects ablated transport activity, while ligand affinity was preserved, allowing for measurements of cytoplasmic pathway accessibility. Although the overall accessibility of the cytoplasmic pathway was markedly reduced under all conditions – and the Na^+^-dependent decrease seen in **Fig. 3A** was not observed (**Fig. 3D**, columns 1 vs 2) – small but significant increases in accessibility were found in the presence of Na^+^ when either 5-HT^+^ or Cl^−^ was added (**Fig. 3D**, columns 2 vs. 3 and 4) with a less significant increase with both 5-HT^+^ and Cl^−^ (**Fig. 3D**, column 5 vs. 4).

### Impact of the Q332E mutation on the molecular network involving Arg104

The Cys277 reactivity of Q332E indicates that the extracellular pathway is more likely to be open than in wild-type SERT, or rather that the intracellular pathway is more likely to be closed; yet, puzzlingly, the mutant retained Cl^−^-dependent changes in cysteine accessibility (**Fig. 3D**). To analyze how a glutamate side chain at position 332 affects the molecular network, we repeated the simulations with Gln332 mutated to glutamate in the presence of 5-HT^+^, Na^+^, and Cl^−^ ions. As envisioned, charged Glu332 interacted more frequently than Gln332 with Arg104; however, this was true only when the pathway was outward-open (**Fig. S6A** left panel; **Fig. S6B**, dark lines) and not for the occluded state (**Fig. S6C**, left panel, **Fig. S6D**, dark lines). Moreover, the Cl^−^ ion placed next to charged Glu332 was inherently unstable and typically left the site within the 500 ns simulation time frame (**Fig. S7B**; green lines). This contrasts starkly with the stable coordination observed in wild-type SERT (**Fig. S7A**; green lines). We attribute the rapid unbinding of the Cl^−^ ion in Q332E to electrostatic repulsion between the Cl^−^ ion and the charge at Glu332, which will be substantial even when Glu332 is rotated away and interacting with Arg104.

We also note that the presence of a Cl^−^ ion profoundly affects the Na^+^ ion at the neighboring Na1 site (**Fig. S7A**, blue; **Fig. S7B**, blue), which unbinds rapidly once the Cl^−^ site is empty (**Fig. S7A**, orange; **Fig. S7B**, orange), in both wild-type and Q332E SERT.

Given the instability of the bound Cl^−^ ion, to study the role of Glu332 in the local network, we repeated the simulations without the Cl^−^ ion. If Gln332 was indeed the primary mediator of Cl^−^binding, we expected that the pathway salt bridge would be less frequently formed in the Glu332 mutant in the absence of Cl^−^ than in wild type. Indeed, when the pathway was open, the Arg104-Glu493 salt bridge was less favored (**Fig. S6A, S6B**, blue) than for wild-type SERT (**Fig. 4A-B**, blue). However, in the occluded state we did not observe a substantial reduction in salt-bridge formation relative to the wild-type protein (blue bars in **Fig. S6C** *cf*. **Fig. 4C**). Thus, the mutation primarily influences the more outward-open states of the transporter in the absence of Cl^−^, stabilizing those conformations and presumably, therefore, inhibiting closure of the extracellular pathway, in line with the lack of measured cytoplasmic accessibility of S277C-Q332E (**Fig. 3D**).

In summary, our simulations suggest, first, that Glu332 reduces the affinity for Cl^−^ in the outward-open state relative to Gln332 (**Fig. S6**), and secondly that in the absence of the ion, the acidic carboxyl side chain in Glu332 in the outward-open conformation strongly attracts Arg104 away from Glu493 (**Fig. S6A**, right), which in turn is expected to increase the energetic barrier to closure of the extracellular pathway (**Fig. S6B**, light blue). In wild-type SERT, by contrast, the Cl^−^ ion appears to act primarily through stabilization of a local network of interactions within the bundle and around Na1 required for formation of the occluded state.

## Discussion

Several observations reported here provide deeper understanding of the unique properties and mechanism of ion-coupled transport of 5-HT^+^. In contrast to two other well-characterized NSS family members, GAT-1 and GlyT1, which transport amino acid neurotransmitters, we show that 5-HT^+^ accumulation by SERT was not affected by imposition of a transmembrane Cl^−^ gradient in either direction (inwardly- or outwardly-directed) in proteoliposomes. This behavior indicates that transmembrane Cl^−^ gradients do not provide a driving force for 5-HT^+^ accumulation, contrary to the expectation if stoichiometric Cl^−^ symport was the basis for chloride dependence.

The inability of Cl^−^ gradients to drive 5-HT accumulation is not likely to reflect any inability of these proteoliposomes to maintain a transmembrane Cl^−^ gradient. Rapid dissipation of that gradient would require compensating flux of a counterion to prevent buildup of a membrane potential. However, gradients of Na^+^, K^+^, and H^+^ were preserved long enough to support substrate accumulation of (**Fig. 2A** and [Ref. (52)]). Moreover, when a K^+^ diffusion potential was imposed in these proteoliposomes with valinomycin, we observed no Cl^—^dependent dissipation of that potential (Fig. 2D). Consequently, we consider rapid dissipation of Cl^−^gradients unlikely to be responsible for the failure of Cl^−^ gradients to affect substrate accumulation.

Taken together, these findings resolve a long-standing ambiguity by showing that chloride is required for SERT function not as a co-transported substrate, but as a permissive or modulatory factor acting within the transport cycle.

Another distinct feature of SERT is that evidence suggests it symports 5-HT^+^ with only 1 Na^+^ ion (6), despite the presence of 2 Na^+^ binding sites in substrate-bound structures (28) and the requirement for two Na^+^ ions for imipramine binding (6). By contrast, GAT-1 and GlyT1 have been shown to symport 2 Na^+^ ions with substrate (20, 53, 54).

This difference in Na⁺ handling has direct implications for whether Cl⁻ can remain stably associated during the transport cycle. Of the two Na^+^ sites, the single Na^+^ ion transported by SERT is most likely to be lost from the Na2 site, given that separation of TM1 and TM8 upon pathway opening severely disrupts the Na2 site (55). Moreover, we expect that site to remain Na^+^-free during the transition to the outward-facing state since intracellular K^+^ binds to the Na2 site to drive substrate accumulation (9, 56), keeping in mind that the K^+^ or H^+^:5-HT^+^ antiport stoichiometry is reported to be 1:1 (8, 57).

What then might explain the differences in Na^+^ stoichiometry from the amino acid transporters? Only the monamine transporters transport amine substrates among all NSS members. Their special property that potentially disfavors Na^+^ dissociation from the Na1 site is that they contain an Asp residue (Asp98 in SERT) that coordinates Na^+^ at Na1 through its side-chain carboxyl group. By contrast, NSS amino acid transporters, including GAT-1 and GlyT1, have a glycine at the position corresponding to SERT Asp98 and use the substrate carboxyl group to coordinate Na^+^ at Na1 (**Fig. 1**) (24, 58). Substrate dissociation from GAT-1 or GlyT1 removes this coordinating group and likely facilitates Na^+^ dissociation from the Na1 site, consistent with their 2:1 Na^+^:amino acid stoichiometry.

Our simulations show a strong synergy of binding between this Na1-site Na^+^ ion and the bound Cl^−^ (**Fig. S7**), consistent with recent simulations of the inward-facing state (56) and the co-dependence of Na^+^ and Cl^−^ affinities in SERT (22). Consequently, we propose that persistent Na1 occupancy in monoamine transporters stabilizes the adjacent Cl^−^ ion, disfavoring its intracellular release together with substrate. Conversely, the loss of the cation from the Na1 site in GAT-1 and GlyT1 would likely destabilize the adjacent Cl^−^ ion, providing an explanation for chloride symport in amino acid transporters.

Even though our results argue against Cl^−^ symport with 5-HT^+^, Cl^−^ is nevertheless required for SERT function (18, 59). This requirement for Cl^−^ resembles the requirement for an acidic group in the bacterial NSS transporters LeuT and TnaT, which contain a Glu or Asp, respectively, at the position in TM7 that Cl^−^-dependent NSS transporters contain a Cl^−^-coordinating serine residue (22, 33, 60) (S372 in SERT, **Fig. 1**).

We show that binding a Cl^−^ ion influences the conformational dynamics of the extracellular side of the transporter by stabilizing the bundle helices, in agreement with simulations examining NaCl binding (61). Importantly, this effect is not mediated solely through formation of the extracellular salt bridge. Surprisingly, the salt bridge spanning the extracellular pathway can be formed independently of chloride binding, even in the outward-open state, despite contrary observations for LeuT in which it is fully dissociated (23). Thus, while Arg104-Glu493 interactions contribute to pathway closure (55), our results indicate that chloride acts more broadly by organizing the extracellular half of the bundle, rather than enforcing a single gating interaction.

To understand why the salt bridge can form in states where the extracellular pathway is open, it is important to note that the carboxyl in SERT is, uniquely, provided by a glutamate side chain, whose longer side chain apparently allows it to reach further across the pathway than the aspartate observed in most other NSS monoamine transporters (**Fig. S8**). Thus, even when the bundle helices including TM1b are angled away from the scaffold, the interaction can form.

SERT also uniquely contains a second glutamate at the neighboring position in TM10 (Glu494; **Fig. S8**), which creates a more electronegative environment in the extracellular pathway sufficient to provide a transient sodium site during binding (61, 63). The presence of Glu494 may explain why SERT-E493C retains transport activity (64), unlike the equivalent GAT-1 mutant (D451C) (65). Analysis of the distances in the MD simulations of SERT revealed that in the presence of the Cl^−^ ion, Arg104 can reach salt-bridging distance to Glu494, albeit only while also contacting Glu493 (**Fig. S3**). An involvement of Glu494 is consistent with the observation that Glu replacements of both Asp451 and Tyr452 in GAT-1 rescues some of the loss of transport seen for D451E alone (65). Together these features indicate that extracellular pathway closure in SERT is governed by a distributed electrostatic network rather than a single salt bridge interaction, distinguishing SERT from other NSS transporters.

The simulation analysis illustrates a significant reduction in helix packing when chloride is not bound, which allows water to enter between the helices (**Fig. 6B**). Given this hydration, and the moderate rates of reactivity measured for Y107C in the presence of NaCl and 5-HT^+^ (**Fig. 3A** middle plot, right bar), we wondered if the solvent accessibility of the side chain at position 107 differs for different conformational states of the wild-type protein simulated here. Indeed, even in the outward-occluded conformation with Cl^−^ bound, the mean and standard deviation of the surface area of Tyr107 accessible to an MTSEA-like probe was 363 ± 8 Å^2^ (**Table 2**). Thus, the side chain retains significant accessibility, despite the nearby extracellular loop 4 (EL4), consistent with the measured rates (**Fig. 3A** middle plot, right bar). Nevertheless, Cl^−^ did influence the reactivity at Y107C (**Fig. 3A**). Extending the surface area analysis to the other states of the transporter (**Table 2**) revealed a subtle increase in solvent accessibility for the Cl^−^-free occluded state, but a significant increase to 382 ± 32 Å^2^ and 410 ± 22 Å^2^ for the outward-open conformation in the presence and absence of the Cl^−^ ion, respectively. Thus, the trend follows that measured using MTSEA: Tyr107 is most reactive in outward-facing conformations, but even in that state, Cl^−^ reduces the accessibility of this region of the transporter. The induced change in the conformational dynamics, however, is more subtle than the major conformational change to the inward-facing state. We posit that the Cl^−^ ion acts to organize and stabilize the packing in the four-helix bundle comprising TM helices 1, 2, 6, and 7 even in outward-facing states of the transporter (**Fig. 7**).

**Fig. 7.**
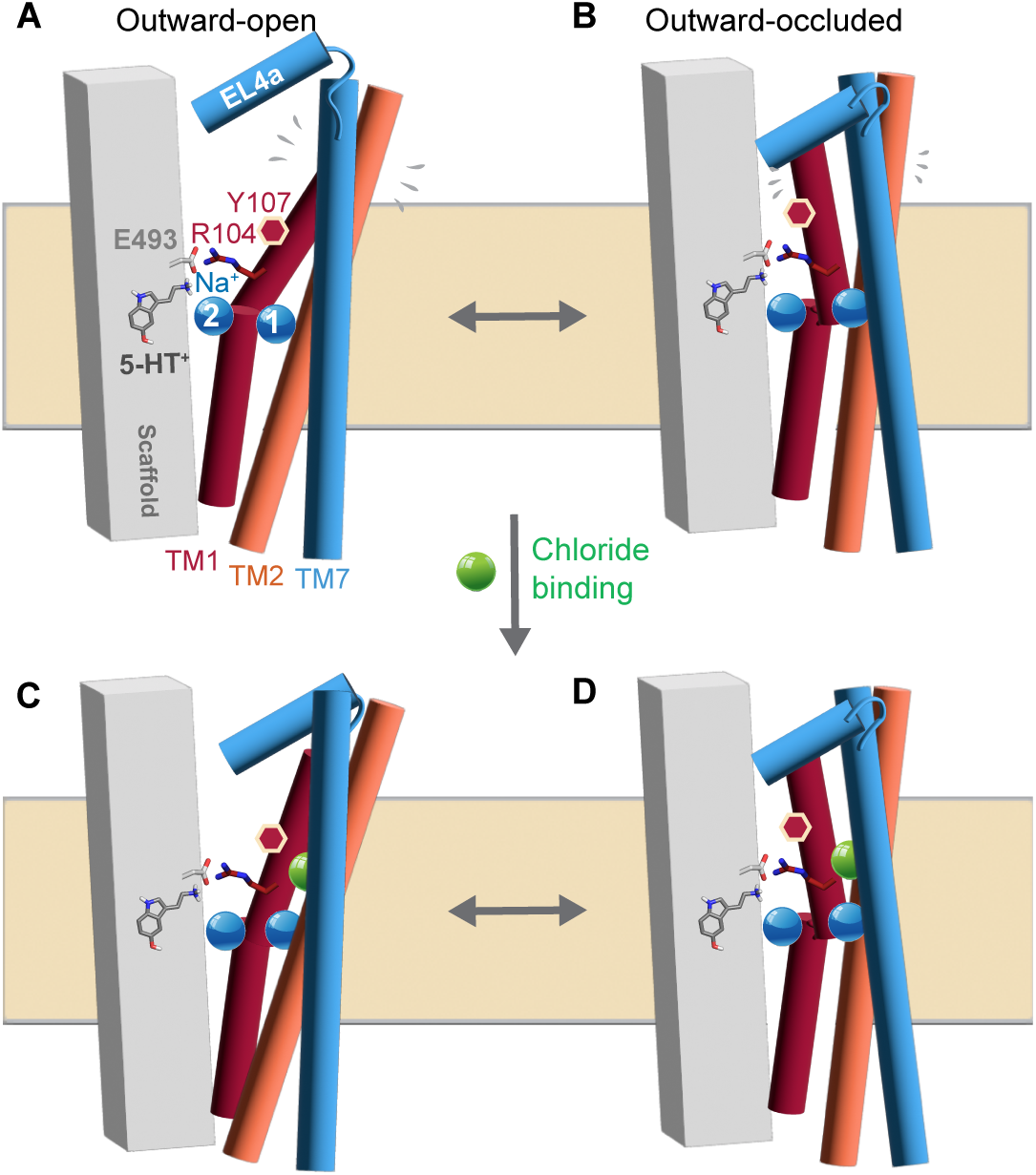
The chloride ion organizes and stabilizes the packing in the four-helix bundle. Schematics of the scaffold region (*grey box*) and the four-helix bundle (TMs 1, 2, 7 are shown as *red, orange, and blue cylinders*, respectively; TM6 is omitted for clarity) highlighting the differences in structural dynamics and the degree of Tyr107 exposure (*red hexagon* in TM1) in the presence **(C, D)** and absence **(A, B)** of chloride (*green sphere*). Serotonin is shown with gray sticks, and the bound Na^+^ ions bound (Na1 and Na2) as blue spheres. O atoms are colored red, N atoms in blue.

No cholesterol molecules were included in our simulations, in part because of the potential for unbinding and the resultant reduced statistical power of the trajectories. Cholesterol in the inner leaflet interacts with SERT at a site near TM5 (66) and impacts substrate affinity and transport kinetics (67), possibly by shifting the transporter equilibrium away from inward-facing states (66). However, the impact of cholesterol on the conformational ensemble of the outward-facing states has yet to be investigated. Further studies are required to assess whether and how cholesterol interactions and the general heterogeneity of the plasma membrane lipid bilayer impact helix packing and ion-dependent conformational dynamics. Given the sensitivity of extracellular helix packing to chloride binding demonstrated here, lipid-mediated modulation of helix interactions—such as that induced by cholesterol—may further tune the same conformational equilibria.

When examining the mutant Q332E, we observed that Cl^−^ rapidly unbinds from the outward-open state, suggesting a very weak affinity. How then can we explain that NaCl modulates the reactivity of S277C experimentally in the Q332E mutant (**Fig. 3B**)? We posit that a small population of protonated Glu332, caused by a shift in *pK_A_* at high Cl^−^ concentrations, will allow a low level of Cl^−^ binding. This explanation remains speculative and is offered as one possible reconciliation of the experimental and simulation results.

A puzzling outcome of recent studies of SERT is that the overall transport cycle is now predicted to involve influx of 2 positive charges (Na^+^:5-HT^+^), and efflux of one positive charge (K^+^), and thus a net electrogenic cycle; this is in contrast to long-held understanding from the impact of ionophores on platelet plasma membrane vesicles that the cycle is electroneutral (7). We consider it possible that a second cation, possibly H^+^, binds in the S1 site and is antiported along with K^+^. However, the nature of this cation is unclear, and further studies are required to resolve this apparent discrepancy and finally establish the stoichiometry of serotonin uptake by hSERT.

To conclude, a combination of uptake and accessibility measurements with simulations finally sheds light on a decades-long question regarding the chloride dependence of SERT, with notable complexity and characteristics distinct from other neurotransmitter transporters. The unique behaviors of SERT may originate from subtle differences in sequence in the extracellular pathway, highlighting the care that must be taken in inferring specific behaviors based on those observed in homologous transporters. Together, our results show that chloride dependence in SERT reflects conformational and ion-binding stabilization within the extracellular bundle rather than chloride symport.

## Materials and Methods

### Reconstitution of SERT Proteoliposomes

Detergent-free reconstitution of SERT proteoliposomes was performed using diisobutylene maleic acid (DIBMA) solubilization as described previously (9) (see **Supporting Information**). The possibility of Cl^−^ leakage was assessed using the fluorescent dye DiS-C3-(5) and by the application of valinomycin, as described in **Supporting Information**.

### Transport Assays with SERT Proteoliposomes

5-HT^+^ uptake was initiated by diluting 10 μL of SERT proteoliposomes (containing 200 ng protein), formed with the indicated internal buffer, into 190 μL of the corresponding external buffer containing 20 nM [^3^H]5-HT (41.3 Ci/mmol, PerkinElmer). After incubation for the indicated time at 22 °C, the reaction was quenched on ice for 1 min and 300 μL of ice-cold external buffer added. Extravesicular [^3^H]5-HT was removed by filtration through glass fiber filters (0.45 μm pore size, Jingteng, China) and the accumulated [^3^H]5-HT in the proteoliposomes was measured by liquid scintillation spectrometry with a Microbeta2 Microplate counter (PerkinElmer). All liposome-based [^3^H]5-HT uptake experiments were performed three times with triplicate measurements in each assay. Nonspecific uptake was measured on liposomes without SERT-DIBMA in parallel.

### Cysteine Accessibility Measurements

Conformational changes were measured using the accessibility of cysteine residues placed in the cytoplasmic (S277C) and extracellular (Y107C) permeation pathways as described previously (10). These constructs retained 70-100% functional activity of the WT protein (68). Two different approaches were used, depending on the pathway of interest. For measurement of extracellular pathway accessibility, [^3^H] 5-HT^+^ uptake measurements were made with intact HeLa cells expressing rat SERT Y107C in the SERT C109A background (69) growing in 96-well culture plates. For cytoplasmic pathway accessibility, measurements were made with membranes prepared from HeLa cells expressing SERT S277C in the X5C background (a SERT mutant in which the five most reactive cysteine were mutated, C15A/C21A/C109A/C357I/C622A) on filters in 96-well filtration plates. A radioligand (β-CIT) binding assay (70) was used with membranes from disrupted cells rather than substrate uptake, because MTSEA does not react with Cys277 in intact cells (71) and consequently does not access the cytoplasmic pathway in intact cells. In both whole cells and membrane fragments, accessibility was measured by the rate of cysteine reactivity with MTSEA, as described previously (10). The MTSEA concentration causing half-maximal inactivation was determined and used together with the time of the reaction (15 min) to calculate pseudo first-order rate constant for cysteine modification, as described previously (4, 40). MTSEA concentrations were calibrated using Ellman’s reagent (5,5′-dithiobis(2-nitrobenzoate)) (72). 5-HT^+^, where added, was present at 10 µM. NMDG^+^ was used to replace Na^+^ and gluconate replaced Cl^−^.

### Molecular dynamics simulations

Simulations of outward-open and outward-occluded conformations of human SERT were prepared according to a protocol described previously (73). Setup and serotonin parameterization details are provided in **Supporting Information (Fig. S9-S10; Tables S3-S4)**.

## Data availability

Input files and representative frames of molecular dynamics simulation trajectories have been deposited in Zenodo [doi: 10.5281/zenodo.14963017], https://zenodo.org, along with serotonin parameter files. All other data are available in the main text or the supplementary materials.

## Author contributions

Conceptualization: EH, GR, LRF

Molecular simulations: EH, JH, AB, RS, EO

Bench experiments: SU, BB, QC, LC

Visualization: EH, JH, AB, LRF

Supervision: EH, YWZ, GR, LRF

Writing—original draft: EH, JH, AB, EO, GR, LRF

Writing—review & editing: EH, EO, YWZ, GR, LRF

Resources, project administration and funding acquisition: YWZ, GR, LRF

## Funding

National Institutes of Health, National Institute of Neurological Disorders and Stroke, grant R01NS102277 (GR)

National Institutes of Health, National Institute of Neurological Disorders and Stroke, Division of Intramural Research NS003139 (LRF)

National Natural Science Foundation of China 32071233 and 32371304 (Y-WZ)

## Acknowledgments

JH was supported by a VDS-PhaNuSpo Mobility Grant of the Vienna Doctoral School of Pharmaceutical, Nutritional and Sport Sciences, University of Vienna, Austria. This work utilized the computational resources of the National Institutes of Health Biowulf high-performance computing cluster and was supported in part by the Intramural Research Program of the NIH. The contributions of the NIH authors were made as part of their official duties as NIH federal employees, are in compliance with agency policy requirements, and are considered Works of the United States Government. However, the findings and conclusions presented in this paper are those of the author(s) and do not necessarily reflect the views of the NIH or the U.S. Department of Health and Human Services. ChatGPT-4o and 5.2 and Claude Haiku were used for suggestions to streamline the language.

## References

1. G. Rudnick, W. Sandtner, Serotonin transport in the 21st century. J Gen Physiol 151, 1248–1264 (2019).

2. C. Fenollar-Ferrer et al., Structure and regulatory interactions of the cytoplasmic terminal domains of serotonin transporter. Biochemistry 53, 5444–5460 (2014).

3. S. Tavoulari et al., Two Na+ Sites Control Conformational Change in a Neurotransmitter Transporter Homolog. J Biol Chem 291, 1456–1471 (2016).

4. Y. W. Zhang, G. Rudnick, The cytoplasmic substrate permeation pathway of serotonin transporter. J Biol Chem 281, 36213–36220 (2006).

5. K. Schicker et al., Unifying concept of serotonin transporter-associated currents. J Biol Chem 287, 438–445 (2012).

6. J. Talvenheimo, H. Fishkes, P. J. Nelson, G. Rudnick, The serotonin transporter-imipramine "receptor". J Biol Chem 258, 6115–6119 (1983).

7. G. Rudnick, P. J. Nelson, Platelet 5-hydroxytryptamine transport, an electroneutral mechanism coupled to potassium. Biochemistry 17, 4739–4742 (1978).

8. P. J. Nelson, G. Rudnick, Coupling between platelet 5-hydroxytryptamine and potassium transport. J Biol Chem 254, 10084–10089 (1979).

9. E. Hellsberg et al., Identification of the potassium-binding site in serotonin transporter. Proc Natl Acad Sci U S A 121, e2319384121 (2024).

10. M. T. Jacobs, Y. W. Zhang, S. D. Campbell, G. Rudnick, Ibogaine, a noncompetitive inhibitor of serotonin transport, acts by stabilizing the cytoplasm-facing state of the transporter. J Biol Chem 282, 29441–29447 (2007).

11. M. W. Quick, Regulating the conducting states of a mammalian serotonin transporter. Neuron 40, 537–549 (2003).

12. I. R. Möller et al., Conformational dynamics of the human serotonin transporter during substrate and drug binding. Nature Communications 10, 1687 (2019).

13. O. Lingjaerde, Jr., Uptake of serotonin in blood platelets in vitro. I. The effects of chloride. Acta Physiol Scand 81, 75–83 (1971).

14. G. Rudnick, Active transport of 5-hydroxytryptamine by plasma membrane vesicles isolated from human blood platelets. J Biol Chem 252, 2170–2174 (1977).

15. M. E. Reith, I. Zimanyi, C. A. O’Reilly, Role of ions and membrane potential in uptake of serotonin into plasma membrane vesicles from mouse brain. Biochem Pharmacol 38, 2091–2097 (1989).

16. D. R. Cool, F. H. Leibach, V. Ganapathy, Modulation of serotonin uptake kinetics by ions and ion gradients in human placental brush-border membrane vesicles. Biochemistry 29, 1818–1822 (1990).

17. H. Gu, S. C. Wall, G. Rudnick, Stable expression of biogenic amine transporters reveals differences in inhibitor sensitivity, kinetics, and ion dependence. J Biol Chem 269, 7124–7130 (1994).

18. P. J. Nelson, G. Rudnick, The role of chloride ion in platelet serotonin transport. J Biol Chem 257, 6151–6155 (1982).

19. P. S. Hasenhuetl, M. Freissmuth, W. Sandtner, Electrogenic Binding of Intracellular Cations Defines a Kinetic Decision Point in the Transport Cycle of the Human Serotonin Transporter. J Biol Chem 291, 25864–25876 (2016).

20. M. J. Roux, S. Supplisson, Neuronal and glial glycine transporters have different stoichiometries. Neuron 25, 373–383 (2000).

21. R. Radian, B. I. Kanner, Stoichiometry of sodium- and chloride-coupled gamma-aminobutyric acid transport by synaptic plasma membrane vesicles isolated from rat brain. Biochemistry 22, 1236–1241 (1983).

22. S. Tavoulari, A. N. Rizwan, L. R. Forrest, G. Rudnick, Reconstructing a chloride-binding site in a bacterial neurotransmitter transporter homologue. J Biol Chem 286, 2834–2842 (2011).

23. Y. W. Zhang et al., Chloride-dependent conformational changes in the GlyT1 glycine transporter. Proc Natl Acad Sci U S A 118 (2021).

24. Y. Wei et al., Transport mechanism and pharmacology of the human GlyT1. Cell 187, 1719–1732 e1714 (2024).

25. B. Lopez-Corcuera, B. I. Kanner, C. Aragon, Reconstitution and partial purification of the sodium and chloride-coupled glycine transporter from rat spinal cord. Biochim Biophys Acta 983, 247–252 (1989).

26. A. Yamashita, S. K. Singh, T. Kawate, Y. Jin, E. Gouaux, Crystal structure of a bacterial homologue of Na^+^/Cl^-^-dependent neurotransmitter transporters. Nature 437, 215–223 (2005).

27. Y. W. Zhang et al., Structural elements required for coupling ion and substrate transport in the neurotransmitter transporter homolog LeuT. Proc Natl Acad Sci U S A 115, E8854–E8862 (2018).

28. D. Yang, E. Gouaux, Illumination of serotonin transporter mechanism and role of the allosteric site. Sci Adv 7, eabl3857 (2021).

29. T. Hu et al., Transport and inhibition mechanisms of the human noradrenaline transporter. Nature 632, 930–937 (2024).

30. J. Tan et al., Molecular basis of human noradrenaline transporter reuptake and inhibition. Nature 632, 921–929 (2024).

31. A. Song, X. Wu, Mechanistic insights of substrate transport and inhibitor binding revealed by high-resolution structures of human norepinephrine transporter. Cell Res 34, 810–813 (2024).

32. Y. Li et al., Dopamine reuptake and inhibitory mechanisms in human dopamine transporter. Nature 632, 686–694 (2024).

33. E. Zomot et al., Mechanism of chloride interaction with neurotransmitter:sodium symporters. Nature 449, 726–730 (2007).

34. H. Krishnamurthy, E. Gouaux, X-ray structures of LeuT in substrate-free outward-open and apo inward-open states. Nature 481, 469–474 (2012).

35. A. Ben-Yona, B. I. Kanner, Functional defects in the external and internal thin gates of the gamma-aminobutyric acid (GABA) transporter GAT-1 can compensate each other. J Biol Chem 288, 4549–4556 (2013).

36. A. V. Pedersen, T. F. Andreassen, C. J. Loland, A conserved salt bridge between transmembrane segments 1 and 10 constitutes an extracellular gate in the dopamine transporter. J Biol Chem 289, 35003–35014 (2014).

37. M. V. Dilworth, H. E. Findlay, P. J. Booth, Detergent-free purification and reconstitution of functional human serotonin transporter (SERT) using diisobutylene maleic acid (DIBMA) copolymer. Biochim Biophys Acta Biomembr 1863, 183602 (2021).

38. A. M. O’Connell, R. E. Koeppe, 2nd, O. S. Andersen, Kinetics of gramicidin channel formation in lipid bilayers: transmembrane monomer association. Science 250, 1256–1259 (1990).

39. P. J. Sims, A. S. Waggoner, C. H. Wang, J. F. Hoffman, Studies on the mechanism by which cyanine dyes measure membrane potential in red blood cells and phosphatidylcholine vesicles. Biochemistry 13, 3315–3330 (1974).

40. S. Tavoulari, L. R. Forrest, G. Rudnick, Fluoxetine (Prozac) binding to serotonin transporter is modulated by chloride and conformational changes. J Neurosci 29, 9635–9643 (2009).

41. I. Singh et al., Structure-based discovery of conformationally selective inhibitors of the serotonin transporter. Cell 186, 2160–2175.e2117 (2023).

42. Y.-W. Zhang et al., Structural elements required for coupling ion and substrate transport in the neurotransmitter transporter homolog LeuT. Proceedings of the National Academy of Sciences 115, 201716870 (2018).

43. S. Tavoulari et al., Two Na+ Sites Control Conformational Change in a Neurotransmitter Transporter Homolog. Journal of Biological Chemistry 291, 1456 1471 (2016).

44. C. Fenollar-Ferrer et al., Structure and regulatory interactions of the cytoplasmic terminal domains of serotonin transporter. Biochemistry 53, 5444–5460 (2014).

45. S. Bulling et al., The mechanistic basis for noncompetitive ibogaine inhibition of serotonin and dopamine transporters. J Biol Chem 287, 18524–18534 (2012).

46. S. Tavoulari, L. R. Forrest, G. Rudnick, Fluoxetine (prozac) binding to serotonin transporter is modulated by chloride and conformational changes. J. Neurosci. 29, 9635–9643 (2009).

47. Y. W. Zhang, G. Rudnick, Cysteine-scanning mutagenesis of serotonin transporter intracellular loop 2 suggests an alpha-helical conformation. J Biol Chem 280, 30807–30813 (2005).

48. J. A. Coleman, E. M. Green, E. Gouaux, X-ray structures and mechanism of the human serotonin transporter. Nature 532, 334–339 (2016).

49. Y. Zhao et al., Single-molecule dynamics of gating in a neurotransmitter transporter homologue. Nature 465, 188–193 (2010).

50. G. Khelashvili et al., Conformational Dynamics on the Extracellular Side of LeuT Controlled by Na+ and K+ Ions and the Protonation State of Glu290. J Biol Chem 291, 19786–19799 (2016).

51. R. Gradisch et al., Occlusion of the human serotonin transporter is mediated by serotonin-induced conformational changes in the bundle domain. Journal of Biological Chemistry 298, 101613 (2022).

52. E. Hellsberg, G. F. Ecker, A. Stary-Weinzinger, L. R. Forrest, A structural model of the human serotonin transporter in an outward-occluded state. PLOS ONE 14, e0217377 (2019).

53. S. Keynan, B. I. Kanner, γ-Aminobutyric acid transport in reconstituted preparations from rat brain: coupled sodium and chloride fluxes Biochemistry 27, 12–17 (1988).

54. M. P. Kavanaugh, J. L. Arriza, R. A. North, S. G. Amara, Electrogenic uptake of gamma-aminobutyric acid by a cloned transporter expressed in Xenopus oocytes. J Biol Chem 267, 22007–22009 (1992).

55. J. A. Coleman et al., Serotonin transporter-ibogaine complexes illuminate mechanisms of inhibition and transport. Nature 569, 141–145 (2019).

56. Z. Zhao, E. Tajkhorshid, Cytosolic K^+^ Binding to the Human Serotonin Transporter. Journal of Chemical Theory and Computation 10.1021/acs.jctc.6c00511 (2026).

57. S. R. Keyes, G. Rudnick, Coupling of transmembrane proton gradients to platelet serotonin transport. J Biol Chem 257, 1172–1176 (1982).

58. A. Zhu et al., Molecular basis for substrate recognition and transport of human GABA transporter GAT1. Nat Struct Mol Biol 30, 1012–1022 (2023).

59. O. Lingjaerde, Uptake of serotonin in blood platelets: Dependence on sodium and chloride, and inhibition by choline. FEBS Lett 3, 103–106 (1969).

60. L. R. Forrest, S. Tavoulari, Y. W. Zhang, G. Rudnick, B. Honig, Identification of a chloride ion binding site in Na+/Cl -dependent transporters. Proc Natl Acad Sci U S A 104, 12761–12766 (2007).

61. D. Szollosi, T. Stockner, Sodium Binding Stabilizes the Outward-Open State of SERT by Limiting Bundle Domain Motions. Cells 11, 255 (2022).

62. J. Pei, B. H. Kim, N. V. Grishin, PROMALS3D: a tool for multiple protein sequence and structure alignments. Nucleic Acids Res 36, 2295–2300 (2008).

63. M. C. Chan, B. Selvam, H. J. Young, E. Procko, D. Shukla, The substrate import mechanism of the human serotonin transporter. Biophys. J. 121, 715–730 (2022).

64. P. C. Keller, 2nd, M. Stephan, H. Glomska, G. Rudnick, Cysteine-scanning mutagenesis of the fifth external loop of serotonin transporter. Biochemistry 43, 8510–8516 (2004).

65. A. Ben-Yona, B. I. Kanner, An acidic amino acid transmembrane helix 10 residue conserved in the neurotransmitter:sodium:symporters is essential for the formation of the extracellular gate of the gamma-aminobutyric acid (GABA) transporter GAT-1. J Biol Chem 287, 7159–7168 (2012).

66. L. Laursen et al., Cholesterol binding to a conserved site modulates the conformation, pharmacology, and transport kinetics of the human serotonin transporter. J Biol Chem 293, 3510–3523 (2018).

67. D. Rudin et al., Cell membrane cholesterol affects serotonin transporter efflux due to altered transporter oligomerization. Mol Psychiatry 10.1038/s41380-025-03201-y (2025).

68. X. Zhang, Y. Xu, Q. Chen, C. Li, Y.-W. Zhang, Control of Conformational Transitions by the Conserved GX9P Motif in the Fifth Transmembrane Domain of Neurotransmitter Sodium Symporters. 10.3390/ijms26073054.

69. Y. Sato, Y. W. Zhang, A. Androutsellis-Theotokis, G. Rudnick, Analysis of transmembrane domain 2 of rat serotonin transporter by cysteine scanning mutagenesis. J Biol Chem 279, 22926–22933 (2004).

70. S. C. Wall, R. B. Innis, G. Rudnick, Binding of the cocaine analog 2 beta-carbomethoxy-3 beta-(4-[125I]iodophenyl)tropane to serotonin and dopamine transporters: different ionic requirements for substrate and 2 beta-carbomethoxy-3 beta-(4-[125I]iodophenyl)tropane binding. Mol Pharmacol 43, 264–270 (1993).

71. A. Androutsellis-Theotokis, G. Rudnick, Accessibility and conformational coupling in serotonin transporter predicted internal domains. J Neurosci 22, 8370–8378 (2002).

72. G. L. Ellman, Tissue sulfhydryl groups. Arch Biochem Biophys 82, 70–77 (1959).

73. X. F. Tan et al., Structure of the Shaker Kv channel and mechanism of slow C-type inactivation. Sci Adv 8, eabm7814 (2022).

74. J. G. Chen, S. Liu-Chen, G. Rudnick, External cysteine residues in the serotonin transporter. Biochemistry 36, 1479–1486 (1997).

