## Supplementary Information for "The Role of Chloride Ions in Serotonin Transport"

PNAS

### Supporting Information for
### The Role of Chloride Ions in Serotonin Transport.

Jiahui Huang, Annika Backer, Stacy Uchendu, Bethlehem Bekele, Chan Li, Qingyang Chen, Esam Orabi, Robyn Stix, Jasper Shide, Yuan-Wei Zhang, Gary Rudnick, Eva Hellsberg, Lucy R. Forrest*

Lucy R. Forrest


**This PDF file includes:**

Supporting text – Materials and Methods
Figures S1 to S10
Tables S1 to S4
SI References

#### Supporting Information Text – Materials and Methods

##### Proteoliposome Experiments

*Reconstitution of SERT Proteoliposomes*. As described previously (1), confluent HEK293T cells stably expressing human SERT wild-type (UniprotKB identifier: P31645) with a Flag epitope at its N- and a 10-His epitope at its C-terminus, respectively, were collected and disrupted by sonication and the cell membrane fraction disrupted in diisobutylene maleic acid (DIBMA) buffer. Native nanodiscs containing SERT were purified on Ni-NTA agarose and Sephadex G-200 size exclusion chromatography to remove imidazole. Lipids (CHS:DOPC:DOPE:DOPG 20:40:25:15) were sonicated and diluted into 20 volumes of the indicated internal buffer, followed by freeze-thaw and extrusion to produce unilamellar liposomes equilibrated with the internal buffer. Purified SERT in native nanodiscs were then reconstituted by addition to the liposomes at a protein/lipid ratio of 1:100 (w/w), vortexed for 1 min, and incubated for 20 min on ice.

*Transport Assays with SERT Proteoliposomes*. 5-$HT^+$ uptake was initiated by diluting 10 µL of SERT proteoliposomes (containing 200 ng protein), formed with the indicated internal buffer, into 190 µL of the corresponding external buffer containing 20 nM [$^3$H]5-HT (41.3 Ci/mmol, PerkinElmer). After incubation for the indicated time at 22 °C, the reaction was quenched on ice for 1 min and 300 µL of ice-cold external buffer added. Extravesicular [$^3$H]5-HT was removed by filtration through glass fiber filters (0.45 µm pore size, Jingteng, China) and the accumulated [$^3$H]5-HT in the proteoliposomes was measured by liquid scintillation spectrometry with a Microbeta2 Microplate counter (PerkinElmer). All liposome-based [$^3$H]5-HT uptake experiments were performed three times with triplicate measurements in each assay. Nonspecific uptake was measured on liposomes without SERT-DIBMA in parallel.

*Testing for $Cl^-$ leaks in reconstituted SERT-DIBMA proteoliposomes*. SERT proteoliposomes (or liposomes free of SERT or DIBMA) were preloaded with an internal buffer (10 mM $Na_2HPO_4$, 130 mM KCl, pH 7.4 or 10 mM $Na_2HPO_4$, 130 mM K-gluconate, pH 7.4) at a lipid concentration of 10 µg/µL and, when present, SERT protein at 0.1 µg/µL. These proteoliposomes were then diluted into an external buffer (10 mM $Na_2HPO_4$ and 130 mM Na- or K-gluconate, pH 7.4) to establish the transmembrane gradient. Fifty microliters of the diluted proteoliposome or liposome suspension (containing 500 µg lipid and 5 µg SERT protein when present) was added to each well of a 96-well plate pre-filled with 200 µl of external buffer, followed by the addition of 1 µL diS-$C_3$-(5) to a final concentration of 1 µM. After shaking for 10 seconds, fluorescence of the cationic dye diS-$C_3$-(5) was measured every 20 seconds for a total duration of 10 minutes using an Infinite 200 Pro microplate reader (Tecan, Männedorf, Switzerland), with excitation and emission wavelengths set to 622 nm and 670 nm, respectively. At 2 min after dye addition, valinomycin was added to each well at a final concentration of 1 µM and at 8 min, Triton X-100 was added to a concentration of 1% (w/v).

##### Cysteine Accessibility Measurements

Conformational changes were measured using the accessibility of cysteine residues placed in the cytoplasmic (S277C) and extracellular (Y107C) permeation pathways as described previously (2). These constructs retained 70-100% functional activity of the WT protein (3). Two different approaches were used, depending on the pathway of interest. For measurement of extracellular pathway accessibility, [$^3$H] 5-HT$^+$ uptake measurements were made with intact HeLa cells expressing rat SERT Y107C in the SERT C109A background (4) growing in 96-well culture plates. For cytoplasmic pathway accessibility, measurements were made with membranes prepared from HeLa cells expressing SERT S277C in the X5C background (a SERT mutant in which the five most reactive cysteine were mutated, C15A/C21A/C109A/C357I/C622A) on filters in 96-well filtration plates. A radioligand (β-CIT) binding assay (5) was used with membranes from disrupted cells rather than substrate uptake, because MTSEA does not react with Cys277 in intact cells (6) and consequently does not access the cytoplasmic pathway in intact cells. In both whole cells and membrane fragments, accessibility was measured by the rate of cysteine reactivity with MTSEA, as described previously (2). The MTSEA concentration causing half-maximal inactivation was determined and used together with the time of the reaction (15 min) to calculate pseudo first-order rate constant for cysteine modification, as described previously (7, 8). MTSEA concentrations were calibrated using Ellman's reagent (5,5′-dithiobis(2-nitrobenzoate)) (9). 5-HT$^+$, where added, was present at 10 μM. NMDG$^+$ was used to replace $Na^+$ and gluconate replaced $Cl^-$.

**Force field parameters for serotonin**

Force field parameters for serotonin were generated using CHARMM General Force Field (CGenFF) parametrization procedures (10, 11). Initial parameters were generated using the CGenFF program website (https://cgenff.silcsbio.com) which assigns parameters by analogy with existing molecular groups and assigns penalty scores to each parameter. For 5-HT$^+$, parameters with penalty scores > 10 were associated with certain ring heavy atoms and with the quaternary N atom (**Fig. S9A**).

Reference data for improving these parameters were generated using quantum chemical calculations. Calculations were performed in the gas phase at the MP2/6-31G(d) level using Gaussian 16. Ab initio geometry optimization revealed two stable conformers for 5-HT$^+$ that differ in the $C_7C_{17}C_{20}N_{23}$ dihedral angle (**Fig. S9C**, blue), leading to a 7.3 kcal/mol difference in energy, as well as distinct dipole moments (**Table S3**). Although penalties assigned to the dihedral angle parameters were low (< 5), their impact on the energy surface for dihedral rotation was tested and compared to that computed with MP2/6-31G(d) (**Fig. S9B-D**). The default CGenFF model correctly predicted the more stable conformer and dipole orientation, though the relative magnitude of the energies differed (**Fig. S9C**, gray, **Table S3**).

The atomic charges of the $CH_2NH_3^+$ fragment were set to those in the ethylammonium ion in CGenFF and the charge on the other $CH_2$ group was set to that of typical CH2 groups (q = –0.18e for C and 0.09e for each H). The atomic charges of other H atoms and of O were left unchanged from the default CGenFF values. We then tested whether these parameters reproduce the interactions of 5-HT$^+$ with water. 5-HT$^+$-water complexes were generated by placing a single $H_2O$ molecule near each of the H-bond donor (C-H, N-H, O-H) and acceptor (O) sites in 5-HT$^+$ (**Fig. S10**). The complexes were set up with a linear H-bond angle between 5-HT$^+$ in its MP2/6-31G(d) optimized geometry and the water molecule with TIP3P geometry ($r_{O\text{-}H}$ = 0.9572 Å, $\theta_{HOH}$ = 104.52°) (12). The interaction distance in these complexes was then optimized at the HF/6-31G(d) level while keeping all other degrees of freedom fixed. The interaction energies of these complexes were calculated without correction for basis set superposition error and together with the H-bond distance were used as target data to adjust the atomic charges on the ring heavy atoms of 5-$HT^+$ (**Table S4**).

The average unsigned error in the default CGenFF parameters relative to the QM values is 12% and 7% for the interaction energies and the H-bond distance, respectively. After adjusting the charges on the ring heavy atoms, the average unsigned error of the interaction energies reduced to 6%, while the H-bond distance error remained at 7%. Importantly, the potential energy surface of the dihedral more closely resembles the QM data after optimization of the charges (**Fig. S9C**, orange).

**Molecular dynamics simulations**

Simulations of two conformations of human SERT were prepared according to a protocol described previously (13). The structure of an outward-occluded conformation was based on a carefully curated hybrid model (14), while an outward-open conformation was based on a structure (Protein Database, PDB identifier 5I71 (15)) including the two reported $Na^+$ ions in sites Na1 and Na2, plus a $Cl^-$ ion assigned the position reported in another structure (PDB identifier 5I6X) (15). Note that this work was initiated before the release of serotonin-bound structures (16). 5-$HT^+$ was added to the S1 binding site according to the position obtained from Induced Fit docking with Schrödinger (14). Force field parameters for 5-$HT^+$ were developed as described below. Residue Glu508 was set to be protonated, based on proximity to Glu136 and the results of our Poisson-Boltzmann calculations to predict the protonated partner of the pair (17) and residues Cys200 and Cys209 were disulfide bridged. Hydrogen atoms in the protein and cavity-bound water were energy minimized for 250 steps using, first, steepest descents and then conjugate gradient protocols. In both cases, the structures were also converted to a coarse-grained representation according to the Martini v2.2 force field (18) and then inserted into a coarse-grained hydrated palmitoylolyeoylphosphatidylcholine (POPC) lipid bilayer at a salt concentration of 150 mM NaCl. Lipids and salt solution were equilibrated around the protein using Gromacs v2018.8 (19) for 50 µs, while constraining the secondary structure of the protein, as described in (20). A single lipid type was used to allow focus on the protein in its simplest model environment and to preclude artefacts due to specific lipid type interactions in different trajectories.

To determine an approximate box size, we analyzed the perturbations of the membrane, finding them to be minimally impacted by the periodic boundary images at a size of 12 x 12 x 11 nm. A representative snapshot with minimal structural deviation from the average equilibrated bilayer was then converted to an all-atom system (21). The protein was replaced by the energy-minimized atomistic sodium- and chloride-bound model and cavities therein were solvated with Dowser (22). Subsequent simulations were carried out with NAMD v2.14 using the CHARMM36m force field (23) and TIP3P waters (12). The periodic box contained ~150,000 atoms with dimensions of ~11 $nm^3$ after coarse-grained equilibration. Two additional wild-type all-atom systems were created by removal of the bound chloride ion which required removal of a bulk sodium ion to balance the net charge of the system.

Structures of Q332E-SERT were derived from the wild-type structure by replacement of the Gln332 side-chain amide group with an O atom. To preserve the net charge, a $Na^+$ ion was added to the bulk solution.

Equilibration of the coarse-grained protein-lipid system was carried out in multiple stages according to (13). In brief, after conversion to the all-atom representation, the lipids, waters, and ions were energy minimized for 100 steps using the conjugate gradient algorithm. To ensure that lipids would not be trapped within protein aromatic rings, pseudo-atoms were placed in the center of each ring during a second 100-step energy minimization. Subsequently, the system was energy minimized for 5,000 steps using the conjugate gradient algorithm with positional restraints on the protein backbone, the nonhydrogen atoms of the side chains, and the O atoms of cavity waters. During a final 500 steps of conjugate gradient minimization, we applied a protein center-of-mass restraint and dihedral restraint on the $\phi$ and $\psi$ and $\chi_1$ angles, with a force constant of 256 kJ/mol. The ions present in the protein binding sites and the Arg104-Glu493 salt bridge were restrained to a minimum number of coordinating interactions using the COLVARS module (24, 25). Specifically, atoms coordinating the ions (Ala96 O, Asn101 $O_{\delta1}$, Ser336 O, Ser336 $O_\gamma$, Asn368 $O_{\delta1}$ in Na1, Gly94 O, Val97 O, Leu434 O, and Ser438 $O_\gamma$ in Na2; and Asn101 $N_{\delta2}$, Tyr121 $O_\eta$, Gln332 $N_{\varepsilon2}$, Ser336 $O_\gamma$, and Ser372 $O_\gamma$ in the $Cl^-$ site) were constrained to ≤2.5 Å from a $Na^+$ ion or ≤3.5 Å from a $Cl^-$ ion, with a force constant of 10 kcal/mol and a minimum coordination number of 2. The extracellular pathway salt bridge comprising Arg104 and Glu493 (between Arg104 $N_\varepsilon$, $N_{\eta1}$ or $N_{\eta2}$ and Glu493 $O_{\delta1}$ or $O_{\delta2}$) was restrained to ≤3.5 Å, with a force constant of 50 kcal/mol and a minimum coordination number of 1. The 5-$HT^+$ present in the S1 site was restrained with distances ≤2.8 Å between Asp98 $O_{\delta2}$ and the cationic nitrogen of 5-$HT^+$, as well as Thr439 $O_{\gamma1}$ and the hydroxyl group of 5-$HT^+$, with a force constant of 200 kcal/mol. The same restraints were applied during six subsequent stages of molecular dynamics (MD) equilibration lasting >204 ns in total, while progressively reducing the force constants. The salt bridge restraint was kept at a force constant of 50 kcal/mol during the first 4 ns, and then progressively reduced by 10 kcal/mol per stage until reaching 10 kcal/mol at equilibration stage 6.

MD simulations were carried out using a 2 fs time step and trajectory output was saved every 4 ps for analyses. A temperature of 298 K was maintained using Langevin Dynamics, and pressure was kept at 1.01325 bar (oscillation time scale = 200 fs; damping scale = 50 fs) using Nosé–Hoover Langevin piston pressure control (26). A cut-off distance of 12 Å with a switching function starting at 10 Å was used for Van der Waals interactions. The particle–mesh Ewald method (27) was used for long-range electrostatic forces. Multiple production runs, each ~0.5 µs long, were carried out without any restraints applied, but with different initial velocities.

*MD Simulation Analysis*. Distance and dihedral angle data were calculated with COLVARS module v2022-05-24 (25) in VMD v1.9.3 (28). The calculated data were visualized and plotted using python matplotlib v3.9.3 (29). Occupancy maps of ions and water were calculated in VMD using the Volmap plugin v1.1 and visualized in VMD and PyMOL version 3.1.3 (Schrödinger Ltd).

**Figures**

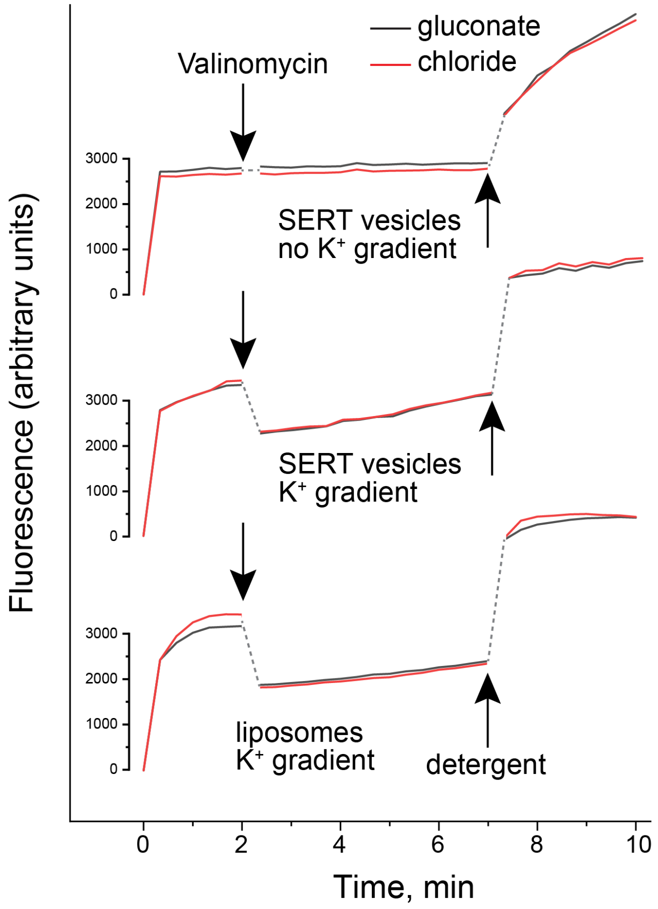


**Fig. S1.**

**Testing the ability of $Cl^-$ to dissipate a valinomycin mediated $K^+$ diffusion potential.** Liposomes prepared in KCl (red traces) or K-gluconate (black traces) were diluted into Na-gluconate to generate a $K^+$ gradient (bottom and center traces) or equimolar K-gluconate as a control (upper trace) and 1 µM diS-$C_3$-(5) was added (0 time). In all traces, fluorescence leveled off within 2 min, although in liposomes containing SERT and DIBMA, a slow steady increase was observed. At 2 min after diS-$C_3$-(5) addition, valinomycin was added to increase $K^+$ permeability in all samples. At 8 min, Triton X-100 was added to disrupt the vesicles. **Bottom and center traces**: liposomes generated in the absence (bottom) or presence (center) of SERT and DIBMA. Addition of valinomycin generated a $K^+$ diffusion potential leading to uptake and quenching of diS-$C_3$-(5), and the presence of $Cl^-$ (red trace) inside the liposomes did not change the response relative to gluconate (black trace), consistent with low $Cl^-$ permeability. **Top trace**: In the absence of a transmembrane $K^+$ gradient, valinomycin addition had no effect on diS-$C_3$-(5) fluorescence in proteoliposomes prepared with SERT and DIBMA.

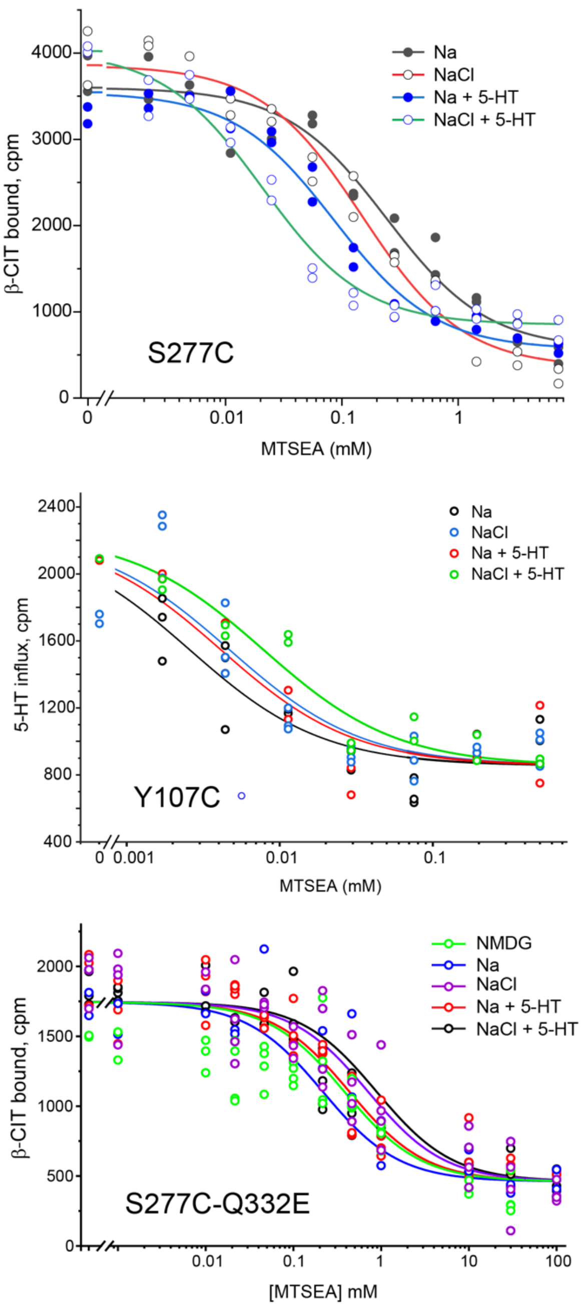


**Fig. S2.**
**Sample curves for reactivity of residues in the SERT pathways.** Titration curves were obtained for accessibility shown as mean ± s.e.m. in the bar charts in Fig. 3A (for S277C), Fig. 3B (for Y107C) and Fig. 3D (for S277C-Q332E). Individual curves display considerable scatter and thus experiments were repeated many times (see **Table S1**).

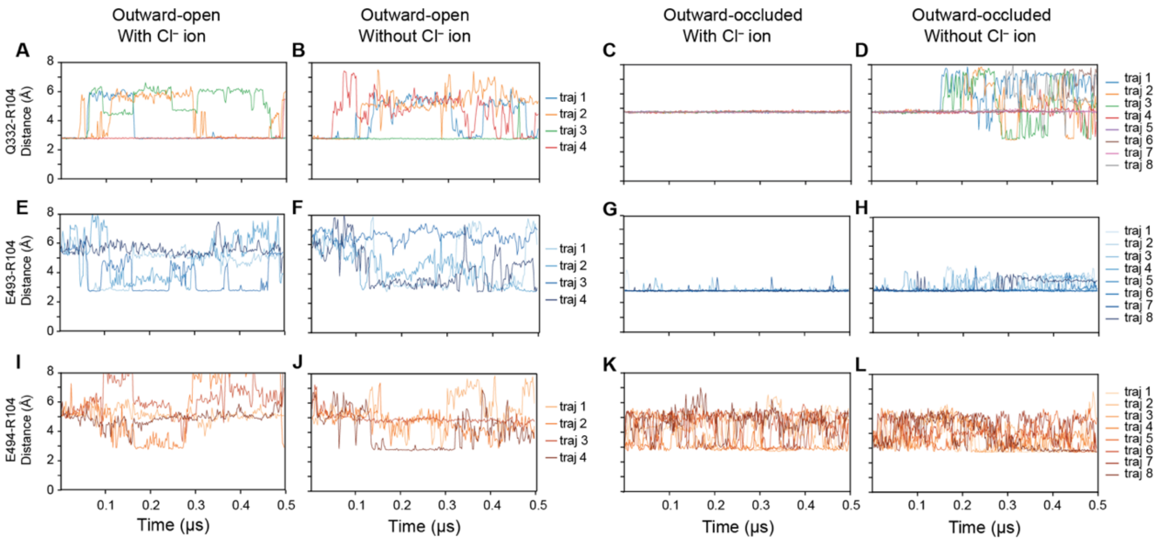


**Fig. S3.**

**Distances between residues in the extracellular salt bridge network as a function of trajectory time in simulations of WT SERT.** Distances were measured as the minimum between polar side-chain N or O atoms in Arg104 and Gln332 **(A-D)**, Glu493 **(E-H)** or Glu494 **(I-L)** and were averaged every 50 datapoints. The transporter was simulated in an outward-open or outward-occluded state, with $Cl^-$ bound or absent, as indicated. Distances < 3.2 Å are considered within range to form an H-bond or salt bridge for the analysis in **Fig. 4** of the main manuscript. Keys indicate the color corresponding to each trajectory of n = 4 or 8 repeats, respectively.

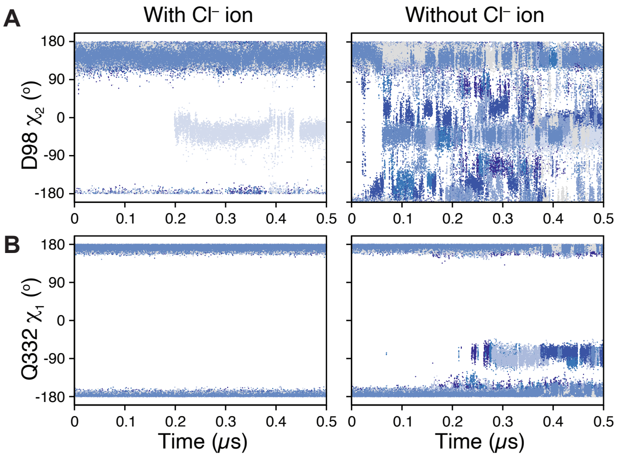


**Fig. S4.**

**Dynamics of ion binding site side chains in the outward-occluded conformation of SERT depend on the presence of the Cl⁻ ion. (A)** Asp98 and **(B)** Gln332 dynamics, measured as the torsion angle of the side chain $\chi_1$ or $\chi_2$ dihedrals, respectively, shown as a function of simulation time for n = 8 trajectories in the presence (*left column*) and absence (*right column*) of a Cl⁻ ion in its binding site. Each trajectory is colored a different shade of blue. Dihedral angles are defined as *trans*, *gauche*(–), *cis* or *gauche*(+) at 180º, 60º, 0º and –60º, respectively. As dihedral angles are continuous, values around ±180º are similar side chain orientations.

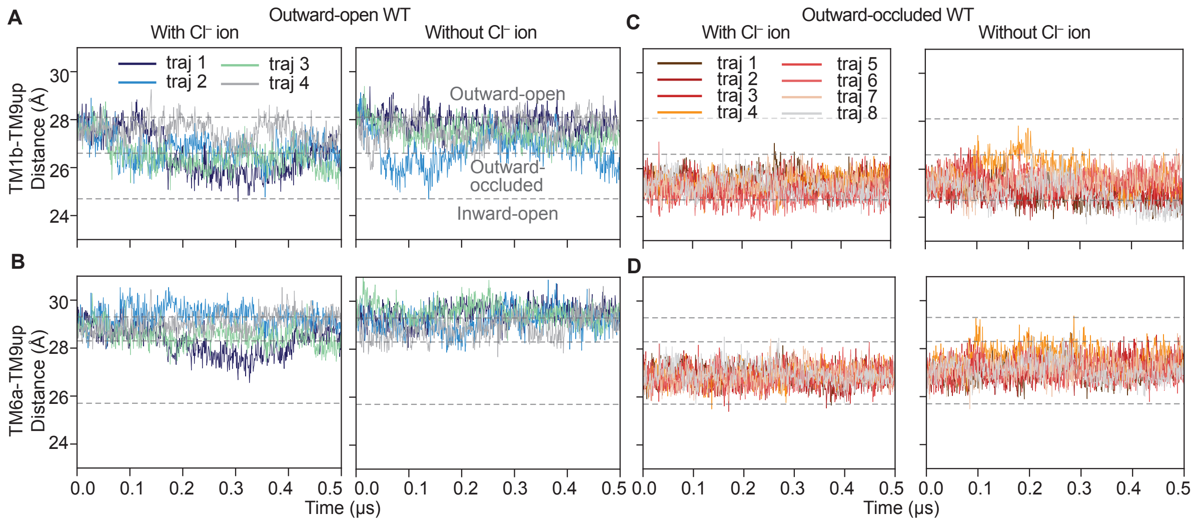


**Fig. S5.**
**Distances between the bundle and scaffold helices during molecular dynamics simulations of wild type SERT**. Simulations of outward-open (**A, B**) or outward-occluded (**C, D**) conformations, both with and without bound $Cl^-$ ion were analyzed. Distances were measured between TM1b (A, C) or TM6a (B, D) in the bundle and the upper segment of TM9 in the scaffold and plotted as a function of simulation time, sampling every 800 ps. Helices were defined as the centers of mass of residues L99 to Q111 (TM1b), residues G324 to L337 (TM6a), and residues F475 to S477 (TM9) following Gradisch et al (30). Red or blue lines indicate different trajectories. Values measured for reported structures in outward-open (PDB identifier 5i71), outward-occluded (7mgw) and inward-open (6dzz) conformations are shown as dashed gray lines.

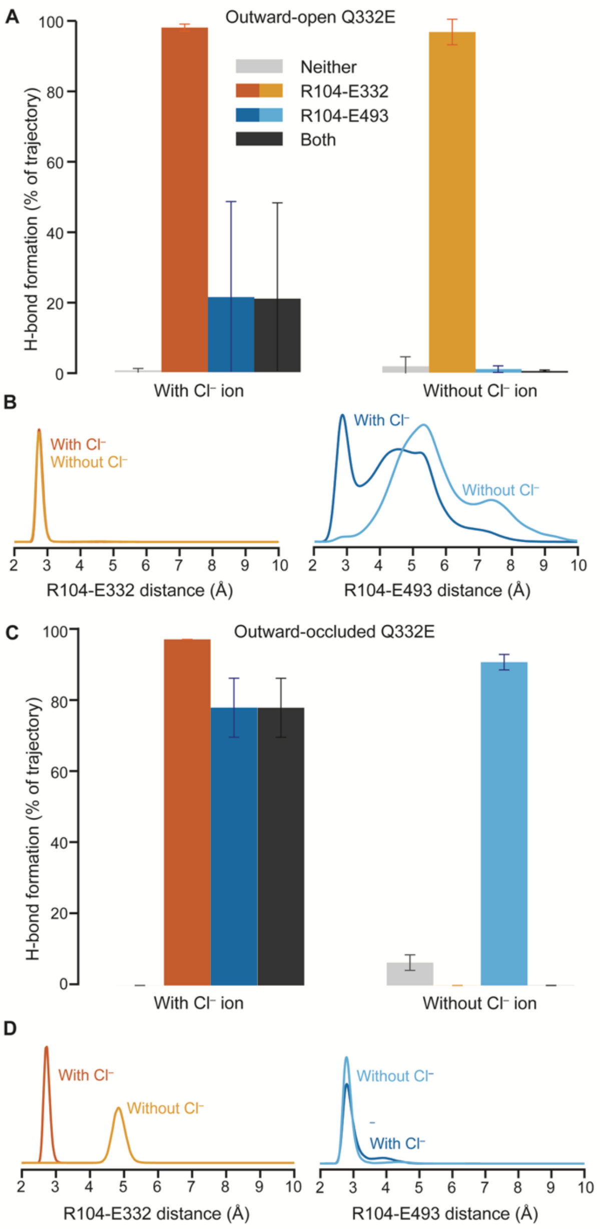


**Fig. S5.**

**Behavior of the interaction network connecting the Cl⁻ binding site to the extracellular pathway salt bridge in the Q332E mutant of SERT.** Simulations were carried out with **(A, B)** outward-open and **(C, D)** outward-occluded conformations for 0.5 μs simulations (n=4 or n=8, respectively). Substrate 5-HT$^+$ and two Na$^+$ ions were included in their respective sites. **(A, C)** Formation of a hydrogen bond (distance <3.2 Å) between any donor atom of Arg104 and any acceptor atom of either the mutated Glu332 (*orange bars*), Glu493 (*blue bars*) or both (*black bars*), either in the presence (*left*) or absence (*right*) of a Cl⁻ ion initially included in its reported binding site. Gray bars represent the fraction of time where neither H-bonds are formed. Error bars reflect standard deviations across repeat trajectories. In (**B, D**) the distance was measured either in the presence (*darker lines*) or the absence (*lighter lines*) of a Cl⁻ ion.

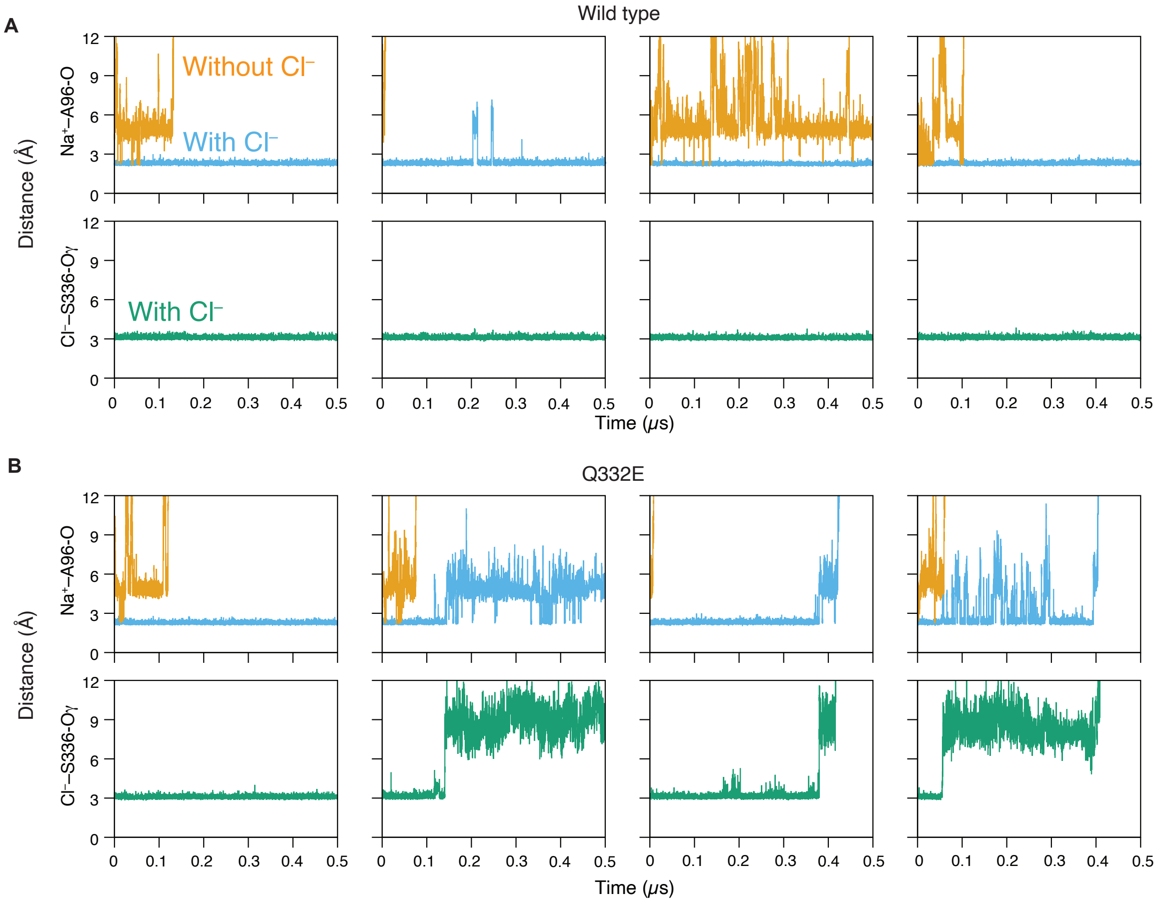


**Fig. S7.**
**Interdependence of $Cl^-$ and $Na^+$ occupancy in simulations of the outward-open conformation of SERT.** Simulations of **(A)** wild type, and **(B)** mutant Q332E were carried out with or without $Cl^-$ bound in at least four independent trajectories (*columns*). The distance of the $Cl^-$ ion, when present, to the sidechain oxygen of Ser336 (*green line*), a representative of the $Cl^-$ site, is compared to the distance of the $Na^+$ ion to the backbone oxygen of Ala96, a representative of the Na1 site, in simulations either with (*cyan*) or without (*orange*) the initially-bound $Cl^-$ ion. Data was sampled every 800 ps.

| Protein | Sequence ID | Sequence Length | Start | TM10 | End |
|---|---|---|---|---|---|
| *hSERT SLC6A4* | PDB 5I71 | 74-615 | 485 | GAYVVKLLEEYAT-GPAVLTVALIEAVAVSWFY | 516 |
| *hDAT SLC6A3* | Q01959 | 1-620 | 468 | GIYVFTLLDHFAA-GTSILFGVLIEAIGVAWFY | 499 |
| *hNET SLC6A2* | P23975 | 1-617 | 465 | GIYVLTLLDTFAA-GTSILFAVLMEAIGVSWFY | 496 |
| *hGAT2 SL6A13* | Q9NSD5 | 1-602 | 439 | GMYVFQLFDYYAASGMCLLFVAIFESLCVAWVY | 471 |
| *hGAT3 SLC6A11* | P48066 | 1-632 | 459 | GMYIFQLFDSYAASGMCLLFVAIFECICIGWVY | 491 |
| *hBGT1 SLC6A12* | P48065 | 1-614 | 444 | GMYIFQLFDYYASSGICLLFLSLFEVVCISWVY | 476 |
| *hTauT SLC6A6* | P31641 | 1-620 | 451 | GMYVFQLFDYYAASGVCLLWVAFFECFVIAWIY | 483 |
| *hCT1 SLC6A8* | P48029 | 1-635 | 466 | GMYVFQLFDYYSASGTTLLWQAFWECVVVAWVY | 498 |
| *hGAT1 SLC6A1* | P30531 | 1-599 | 443 | GIYVFKLFDYYSASGMSLLFLVFFECVSISWFY | 475 |
| *hGlyT2 SLC6A5* | Q9Y345 | 1-797 | 625 | GIYMFQLVDTYAA-SYALVIIAIFELVGISYVY | 656 |
| *hPROT SLC6A7* | Q99884 | 1-636 | 446 | GMYWLVLLDDYSA-SFGLMVVVITTCLAVTRVY | 477 |
| *hGlyT1 SLC6A9* | P48067 | 1-706 | 520 | GIYWLLLMDNYAA-SFSLVVISCIMCVAIMYIY | 551 |
| *hATB0+ SLC6A14* | Q9UN76 | 1-642 | 470 | GIYWVHLIDHFCA-GWGILIAAILELVGIIWIY | 501 |
| *hB0AT1 SLC6A19* | Q695T7 | 1-634 | 478 | GQYWLSLLDSYAG-SIPLLIIAFCEMFSVVYVY | 509 |
| *hB0AT3 SLC6A18* | Q96N87 | 1-628 | 464 | GNYWLEIFDNFAA-SPNLLMLAFLEVVGVVYVY | 495 |
| *hSIT1 SLC6A20* | Q9NP91 | 1-592 | 453 | GNYWFDIFNDYAA-TLSLLLIVLVETIAVCYVY | 484 |
| *hB0AT2 SLC6A15* | Q9H2J7 | 1-730 | 517 | GNYFVTMFDDYSA-TLPLLIVVILENIAVCFVY | 548 |
| *hNTT4 SLC6A17* | Q9H1V8 | 1-727 | 516 | GNYFVTMFDDYSA-TLPLTLIVILENIAVAWIY | 547 |
| *hNTT5 SLC6A16* | Q9GZN6 | 1-736 | 558 | GSYFIRLLSDYWI-VFPIIVVVVFETMAVSWAY | 589 |

**Fig. S8.**

**Multiple Sequence Alignment of the 19 SLC6 transporters.** The full UniProt sequences and SERT structure PDB ID 5I71 were aligned using Promals3D (31). The figure shows the aligned residues of TM10, using SERT as a reference. Glu493 and Glu494 in SERT are highlighted together with the corresponding positions in the other SLC6 members by a purple box. Fully conserved residues are highlighted on an orange background.

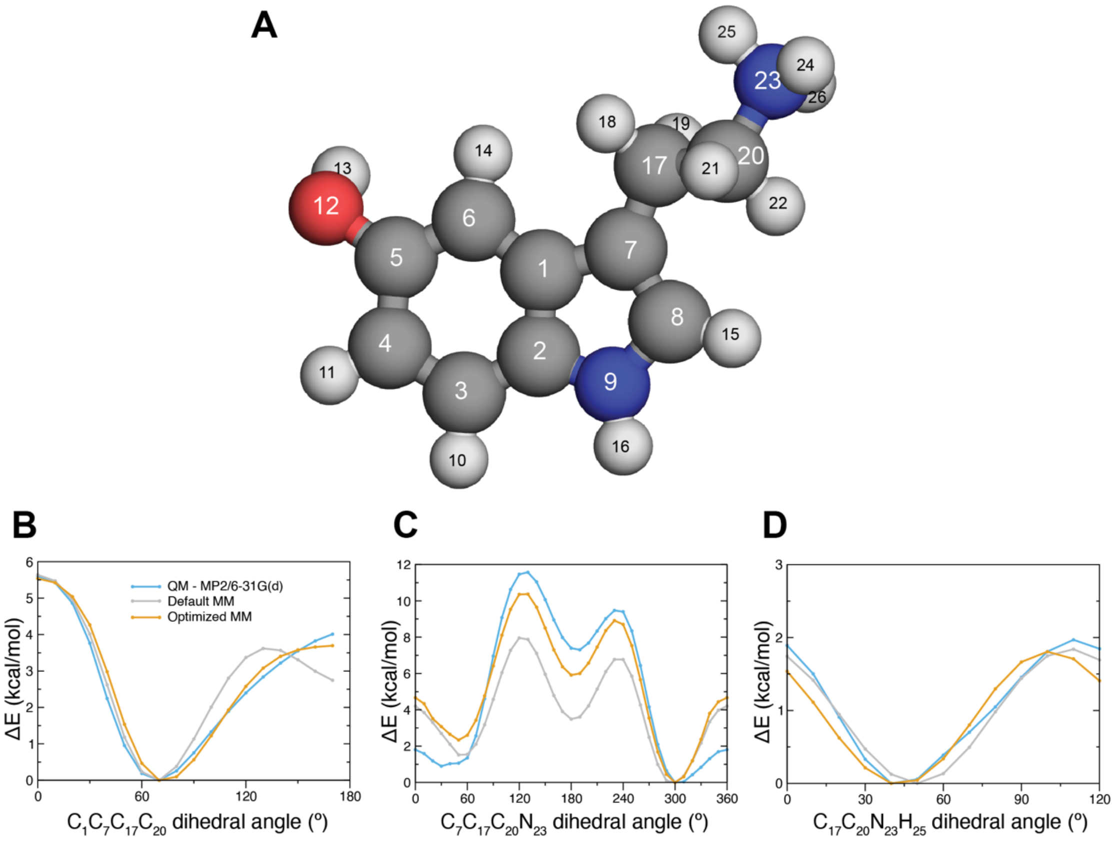


**Fig. S9.**
**Parameterization of the 5-HT$^+$ force field**. (A) Atomic structure of 5-HT$^+$, indicating atom labels used. (B-D) Potential energy curves calculated with MP2/6-31G(d) (*blue*) and with the default (*gray*) and optimized (*orange*) CGenFF, for various dihedrals and plotted as the energy relative to that of the equilibrium angle. The dihedral angles examined were: (B) the $C_1C_7C_{17}C_{20}$ dihedral angle between 0° and 180°, (C) the $C_7C_{17}C_{20}N_{23}$ dihedral angle between 0° and 360°, and (D) the $C_{17}C_{20}N_{23}H_{25}$ dihedral angle between 0° and 120°. Note that, the parameters of the torsions in $C_7C_{17}C_{20}N_{23}$ (C) and $C_{17}C_{20}N_{23}H_{25}$ (D) were kept unmodified; nevertheless, the optimized model produces a superior potential energy surface for $C_7C_{17}C_{20}N_{23}$ due to updated atomic charges.

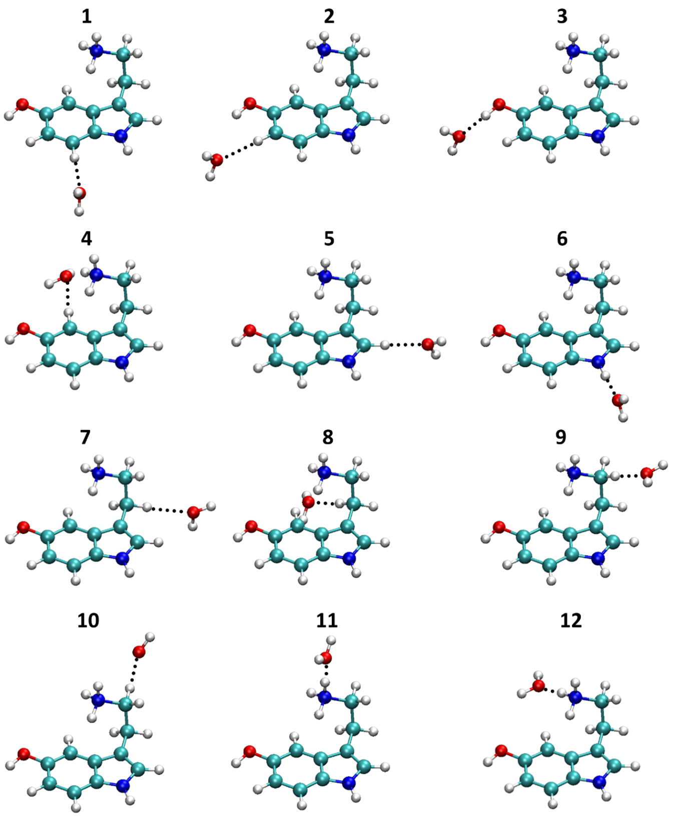


**Fig. S10.**
**Ab initio optimized geometries of 5-HT$^{+}$-water complexes.** These complexes were used in the calibration of an optimized molecular mechanics forcefield for 5-HT$^{+}$.

#### Supplementary Tables

#### Table S1. Data underlying bar charts in Figure 3

**Figure 3A**

| | NMDG | Na | Na + 5-HT | NaCl | NaCl + 5-HT |
|---|---|---|---|---|---|
| n | 22 | 21 | 13 | 21 | 13 |
| Mean | 17.6 | 5.77 | 9.51 | 9.32 | 30.5 |
| SD | 12.0 | 2.73 | 4.29 | 4.34 | 13.0 |
| SEM | 2.56 | 0.57 | 1.19 | 0.95 | 3.61 |

**Figure 3B**

| | Na | Na + 5-HT | NaCl | NaCl + 5-HT |
|---|---|---|---|---|
| n | 10 | 4 | 3 | 3 |
| Mean | 194 | 109 | 96.8 | 63.6 |
| SD | 53.4 | 22.7 | 16.5 | 8.4 |
| SEM | 16.9 | 11.3 | 9.5 | 4.8 |

**Figure 3D**

| | NMDG | Na | Na + 5-HT | NaCl | NaCl + 5-HT |
|---|---|---|---|---|---|
| n | 11 | 11 | 7 | 5 | 9 |
| Mean | 0.51 | 0.73 | 1.41 | 1.94 | 2.55 |
| SD | 0.23 | 0.62 | 0.78 | 0.70 | 1.92 |
| SEM | 0.07 | 0.19 | 0.30 | 0.31 | 0.64 |

**Table S2. Surface accessibility of Tyr107 during MD simulations of SERT**

| Conformation | $Cl^-$ bound | Mean (Å$^2$) | SD | Num of trajectories |
|---|---|---|---|---|
| Outward-open | – | 410 | 22 | 4 |
| | + | 382 | 32 | 4 |
| Outward-occluded | – | 369 | 12 | 8 |
| | + | 363 | 8 | 8 |

Outward-facing state simulations are with bound 5-$HT^+$ and $Na^+$. The surface is computed by rolling a large probe (roughly approximating a methyl-thiosulfonate reagent) with radius 3 Å over the surface of all atoms at each time step. Mean surface area was computed over all frames in all trajectories.

#### Table S3. Properties of the two most stable 5-HT$^+$ conformers obtained from quantum chemical calculations

| | Dipole moment (D) | | D$E$ = $E$(1) – $E$(2), kcal/mol |
|---|---|---|---|
| | Conformer 1 | Conformer 2 | |
| MP2/6-31G(d) | 13.48 | 7.91 | 7.3 |
| Default CGenFF | 14.76 | 9.44 | 3.6 |
| Optimized CGenFF | 12.01 | 7.86 | 5.7 |

#### Table S4. Properties of 5-HT$^+$-$H_2O$ complexes obtained from quantum chemical calculations

| 5-HT$^+$-$H_2O$ Complex | HF/6-31G(d) | | Default CGenFF | | Optimized CGenFF | |
|---|---|---|---|---|---|---|
| | $r_{H...O}$, Å | $E$ (kcal/mol) | $r_{H...O}$, Å | $E$ (kcal/mol) | $r_{H...O}$, Å | $E$ (kcal/mol) |
| 1 | 2.33 | -5.8 | 2.57 | -4.5 | 2.53 | -5.4 |
| 2 | 2.31 | -6.1 | 2.51 | -5.1 | 2.50 | -6.1 |
| 3 | 1.90 | -11.3 | 1.84 | -10.0 | 1.82 | -11.1 |
| 4 | 2.29 | -11.0 | 2.22 | -9.6 | 2.23 | -10.5 |
| 5 | 2.26 | -7.0 | 2.19 | -7.0 | 2.21 | -6.6 |
| 6 | 1.95 | -10.5 | 1.84 | -9.1 | 1.82 | -10.2 |
| 7 | 2.33 | -6.6 | 2.55 | -6.0 | 2.58 | -5.2 |
| 8 | 2.32 | -7.8 | 2.51 | -9.1 | 2.51 | -8.0 |
| 9 | 2.26 | -8.1 | 2.49 | -8.1 | 2.51 | -7.5 |
| 10 | 2.23 | -8.5 | 2.48 | -8.4 | 2.48 | -8.1 |
| 11 | 1.84 | -16.2 | 1.72 | -18.3 | 1.72 | -17.9 |
| 12 | 1.85 | -17.2 | 1.73 | -19.9 | 1.72 | -19.0 |

Interaction H-bond distances ($r$, in Å) and interaction energies ($E$, in kcal/mol) for 5-HT$^+$-water complexes (structures 1-12 in **Fig. S10**). QM data are compared with MM values calculated with default CGenFF and with the optimized forcefield.

#### SI References


1. E. Hellsberg *et al.*, Identification of the potassium-binding site in serotonin transporter. *Proc Natl Acad Sci U S A* **121**, e2319384121 (2024).
2. M. T. Jacobs, Y. W. Zhang, S. D. Campbell, G. Rudnick, Ibogaine, a noncompetitive inhibitor of serotonin transport, acts by stabilizing the cytoplasm-facing state of the transporter. *J Biol Chem* **282**, 29441-29447 (2007).
3. X. Zhang, Y. Xu, Q. Chen, C. Li, Y.-W. Zhang, Control of Conformational Transitions by the Conserved GX9P Motif in the Fifth Transmembrane Domain of Neurotransmitter Sodium Symporters. http://dx.doi.org/10.3390/ijms26073054.
4. Y. Sato, Y. W. Zhang, A. Androutsellis-Theotokis, G. Rudnick, Analysis of transmembrane domain 2 of rat serotonin transporter by cysteine scanning mutagenesis. *J Biol Chem* **279**, 22926-22933 (2004).
5. S. C. Wall, R. B. Innis, G. Rudnick, Binding of the cocaine analog 2 beta-carbomethoxy-3 beta-(4- [125I]iodophenyl)tropane to serotonin and dopamine transporters: different ionic requirements for substrate and 2 beta-carbomethoxy-3 beta-(4-[125I]iodophenyl)tropane binding. *Mol Pharmacol* **43**, 264-270 (1993).
6. M. Holmgren, Y. Liu, Y. Xu, G. Yellen, On the use of thiol-modifying agents to determine channel topology. *Neuropharmacology* **35**, 797-804 (1996).
7. Y. W. Zhang, G. Rudnick, The cytoplasmic substrate permeation pathway of serotonin transporter. *J Biol Chem* **281**, 36213-36220 (2006).
8. S. Tavoulari, L. R. Forrest, G. Rudnick, Fluoxetine (Prozac) binding to serotonin transporter is modulated by chloride and conformational changes. *J Neurosci* **29**, 9635-9643 (2009).
9. G. L. Ellman, Tissue sulfhydryl groups. *Arch Biochem Biophys* **82**, 70-77 (1959).
10. K. Vanommeslaeghe *et al.*, CHARMM general force field: A force field for drug-like molecules compatible with the CHARMM all-atom additive biological force fields. *J Comput Chem* **31**, 671-690 (2010).
11. K. Vanommeslaeghe, A. D. MacKerell, Jr., Automation of the CHARMM General Force Field (CGenFF) I: bond perception and atom typing. *J Chem Inf Model* **52**, 3144-3154 (2012).
12. W. L. Jorgensen, J. Chandrasekhar, J. D. Madura, R. W. Impey, M. L. Klein, Comparison of Simple Potential Functions for Simulating Liquid Water. *Journal of Chemical Physics* **79**, 926-935 (1983).
13. X. F. Tan *et al.*, Structure of the Shaker Kv channel and mechanism of slow C-type inactivation. *Sci Adv* **8**, eabm7814 (2022).
14. E. Hellsberg, G. F. Ecker, A. Stary-Weinzinger, L. R. Forrest, A structural model of the human serotonin transporter in an outward-occluded state. *PLOS ONE* **14**, e0217377 (2019).
15. J. A. Coleman, E. M. Green, E. Gouaux, X-ray structures and mechanism of the human serotonin transporter. *Nature* **532**, 334-339 (2016).
16. D. Yang, E. Gouaux, Illumination of serotonin transporter mechanism and role of the allosteric site. *Sci Adv* **7**, eabl3857 (2021).

17. V. M. Korkhov, M. Holy, M. Freissmuth, H. H. Sitte, The conserved glutamate (Glu136) in transmembrane domain 2 of the serotonin transporter is required for the conformational switch in the transport cycle. *J Biol Chem* **281**, 13439-13448 (2006).
18. S. J. Marrink, H. J. Risselada, S. Yefimov, D. P. Tieleman, A. H. de Vries, The MARTINI force field: coarse grained model for biomolecular simulations. *J Phys Chem B* **111**, 7812-7824 (2007).
19. H. Bekker, Berendsen, H., Dijkstra, E.J., Achterop, S., Drunen, R., van der Spoel, D., Sijbers, A., Keegstra, H., Reitsma, B., Renardus, M.K.R. (1993) Gromacs - A parallel computer for molecular dynamics simulations. in *4th International Conference on Computational Physics*, ed R. A. DeGroot, Nadrchal, J. (World Scientific Publishing, Czech Republic), pp 252-256.
20. R. Stix *et al.*, Eukaryotic Kv channel Shaker inactivates through selectivity filter dilation rather than collapse. *Sci Adv* **9**, eadj5539 (2023).
21. T. A. Wassenaar, K. Pluhackova, R. A. Bockmann, S. J. Marrink, D. P. Tieleman, Going Backward: A Flexible Geometric Approach to Reverse Transformation from Coarse Grained to Atomistic Models. *J Chem Theory Comput* **10**, 676-690 (2014).
22. L. Zhang, J. Hermans, Hydrophilicity of cavities in proteins. *Proteins* **24**, 433-438 (1996).
23. J. Huang *et al.*, CHARMM36m: an improved force field for folded and intrinsically disordered proteins. *Nat Methods* **14**, 71-73 (2017).
24. G. Fiorin, M. L. Klein, J. Hénin, Using collective variables to drive molecular dynamics simulations. *Mol Phys* **111**, 3345-3362 (2013).
25. G. Fiorin *et al.*, Expanded Functionality and Portability for the Colvars Library. *J Phys Chem B* **128**, 11108-11123 (2024).
26. S. E. Feller, Y. H. Zhang, R. W. Pastor, B. R. Brooks, Constant-Pressure Molecular-Dynamics Simulation - the Langevin Piston Method. *Journal of Chemical Physics* **103**, 4613-4621 (1995).
27. T. Darden, D. York, L. Pedersen, Particle Mesh Ewald - an N.Log(N) Method for Ewald Sums in Large Systems. *Journal of Chemical Physics* **98**, 10089-10092 (1993).
28. W. Humphrey, A. Dalke, K. Schulten, VMD: visual molecular dynamics. *J Mol Graph* **14**, 33-38, 27-38 (1996).
29. J. D. Hunter, Matplotlib: A 2D graphics environment. *Comput. Sci. Eng.* **9**, 90-95 (2007).
30. R. Gradisch *et al.*, Occlusion of the human serotonin transporter is mediated by serotonin-induced conformational changes in the bundle domain. *Journal of Biological Chemistry* **298**, 101613 (2022).
31. J. Pei, B. H. Kim, N. V. Grishin, PROMALS3D: a tool for multiple protein sequence and structure alignments. *Nucleic Acids Res* **36**, 2295-2300 (2008).
